# Reconstitution and Expression of *mcy* Gene Cluster in The Model Cyanobacterium *Synechococcus* 7942 Reveals a Roll of MC-LR in Cell Division

**DOI:** 10.1101/2022.02.17.480855

**Authors:** Yanli Zheng, Chunling Xue, Hui Chen, Anqi Jia, Liang Zhao, Junli Zhang, Lixin Zhang, Qiang Wang

## Abstract

Cyanobacterial blooms pose a serious threat to public health due to the presence of cyanotoxins. The most common cyanotoxins, microcystins (MCs), can cause acute poisoning at high concentrations and hepatocellular carcinoma following chronic exposure. Among all MC variants, MC-LR produced by *Microcystis aeruginosa* PCC 7806 is the most common toxic MC. Although the biosynthetic pathway for MC-LR has been proposed, experimental support of this pathway is lacking. In an effort to experimentally validate this pathway, we expressed the 55 kb microcystin biosynthetic gene cluster (*mcy* cluster) (*mcyA–J*) and produced MC-LR in the model cyanobacterium *Synechococcus* 7942. We designed and constructed the strong bidirectional promoter *biPpsbA_2_* between *mcyA* and *mcyD*, reassembled the *mcy* cluster in yeast by transformation-associated recombination (TAR cloning), transformed the gene cluster into the *NSII* site of *Synechococcus* 7942, and successfully expressed MC-LR at a level of 0.006–0.018 fg cell^−1^ day^−1^. The expression of MC-LR led to abnormal cell division and the filamentation of *Synechococcus* 7942 cells, further analysis proved a role of MC-LR in functional assembly of the cell division protein FtsZ, by competing its GTP binding site. These results represent the first synthetic biological expression of the *mcy* cluster and the autotrophic production of MC-LR in a photosynthetic model organism, which lays the foundation for resolving the MC biosynthesis pathway. The suggested role of MC-LR in cell division reveals a mechanism of how blooming cyanobacteria gain a competitive edge over their non-blooming counterparts.

**Highlights:**

1. We expressed the 55 kb *mcy* cluster and produced MC-LR in the model cyanobacterium *Synechococcus* 7942.
2. This is the first realized production of MC in the model non-toxin-production cyanobacteria from CO_2_ by photosynthesis.
3. Compared with the self-replicating plasmid, the recombination of the *mcy* cluster into the genome of *Synechococcus* 7942 is more suitable for the heterologous production of microcystin.
4. MC-LR inhibits cell division by irreversibly competing the GTP binding domain of the cell division protein FtsZ.
5. The newly discovered effect of MC-LR on cell division reveals a mechanism of how blooming cyanobacteria gain competitive edge over their non-blooming counterparts.

## Introduction

Cyanobacterial blooms are a major environmental problem worldwide, posing a serious threat to public health. Blooms can be strengthened by eutrophication and by a combination of several environmental factors such as high water temperature, high salinity, water stagnation, and light intensity (Vasconcelos 2006; Merel et al. 2013). These blooms threaten the health of humans and animals such as fish, crustaceans, mollusks, zooplankton, and birds due to the production of cyanobacterial toxins (Zhou et al. 2021). The most common cyanotoxins, MCs are primarily produced by the *Microcystis* (Zhou et al. 2021). However, other genera such as *Anabaena, Planktothrix, Anabaenopsis, Oscillatoria*, and *Nostoc* can also generate MCs (Hitzfeld, Hoger, and Dietrich 2000; van Apeldoorn et al. 2007; Zhou et al. 2021). MCs are highly hepatotoxic to animals. An emergency case occurred in 2014 in Toledo, OH (Lake Erie, US) that caused authorities to shut down the city’s water supply due to elevated levels of MCs (Cha and Stow 2015). In the most serious case, at least 50 people died of acute liver poisoning due to the use of microcystin-contaminated water in a hemodialysis unit in Caruaru, Brazil (Ochimsen et al. 1998).

MCs have a cyclic hepta-peptide structure (Adda-D-Glu-Mdha-D-Ala-L-X-D-MeAsp-L-Z-) in which X and Z represent variable amino acids, Adda is (2S,3S,8S,9S)-3-amino-9-methoxy-2,6,8-trimethyl-10-phenyldeca-4,6-dienoic acid, Mdha is N-methyldehydroalanine, and D-MeAsp is D-erythro-β-methyl-aspartic acid (Botes et al. 1984; Rinehart, Namikoshi, and Choi 1994; Zhou et al. 2021). The Adda moiety, which is present in all variants, is essential for MC toxicity (Campos and Vasconcelos 2010). The different L-amino acid residues at positions 2 (X) and 4 (Z) account for differences in the toxicokinetic and toxicodynamic properties of MCs (Rinehart, Namikoshi, and Choi 1994). There are many MC variants; to date, more than 270 MC isoforms have been identified (Santori et al. 2020; Massey et al. 2020; Bouaicha et al. 2019; Carmichael 2018). For example, the most common MC congener, MC-LR, contains leucine (L) at position 2 and arginine (R) at position 4.

The *mcy* cluster in *Microcystis aeruginosa* PCC 7806 (*M. aeruginosa* PCC 7806) spans a 55 kb region containing 10 genes (*mcyA-J*). MC biosynthesis is catalyzed by a hybrid enzyme comprising polyketide synthase (PKS)/nonribosomal peptide synthetase (NRPS) (Dittmann 1997; Moore 1991; Arment 1996). A bidirectional promoter is present between *mcyA* and *mcyD*, and the *mcy* cluster is divided into two operons (*mcyA-C* and *mcyD-J*) (Kaebernick et al. 2002). McyA-C are NRPSs, McyD is a PKS, McyE and McyG are PKS/NRPS hybrids, McyF is involved in epimerization, McyI is involved in dehydration, McyJ is an O-methyltransferase, and McyH is a putative toxin transporter (Tillett et al. 2000; Sielaff et al. 2003; Rantala et al. 2006; Pearson, Barrow, and Neilan 2007; Christiansen et al. 2003; Cullen et al. 2019; Pearson et al. 2004).

The microcystin biosynthesis pathway was first proposed by Elke Dittmann and co-workers in 2000 (Tillett et al. 2000). This pathway involves 48 separate biosynthetic reactions, including adenylation, condensation, methylation, reduction, dehydrogenation, and dehydration. The pathway begins with the synthesis of the amino acid Adda. The starting substrate was presumed to be phenylacetate to match the number of carbon atoms in the side chain of Adda. However, Moore et al. and Hicks et al. successively reported that phenylpropanoids but not phenylacetate are the substrate for the synthesis of Adda (Moore 1991; Hicks et al. 2006). However, there would be an extra carbon atom in the side chain of Adda if the starter unit were phenylpropanoids. Therefore, phenylpropanoids, the initial substrate of microcystin biosynthesis, may undergo decarboxylation during the first hydrocarbon chain extension reaction, however, the specific reaction process and some intermediate steps in microcystin biosynthesis pathway are Unclear. For example, D-glutamic acid is incorporated into the sixth position of MCs, but the enzymes involved in this process have not been identified (Cullen et al. 2019; Sielaff et al. 2003).

To reassemble the *mcy* cluster, Liu and co-workers constructed a genomic fosmid library of *M. aeruginosa* PCC 7806. Two microcystin variants (MC-LR and [D-Asp^3^]-MC-LR) were synthesized in *E. coli* (Liu et al. 2017). The authors synthesized the pFos-mcy plasmid carrying the *mcy* cluster by fosmid library and used it to transform *E. coli* GB05-MtaA cells. The native promoter located between *mcyA* and *mcyD* was used to drive *mcy* expression in the pFos-mcy plasmid; however, no microcystin was produced by the *E. coli* cells. The authors then constructed the bidirectional inducible promoter *PtetO* by Gibson cloning to replace the native promoter, followed by transformation into *E. coli.* The *mcy* cluster was successfully transcribed, and [D-Asp^3^]-MC-LR was produced. MC-LR was also produced by the *E. coli* cells following the addition of the amino acid β-methyl-aspartic acid into the culture medium due to the absence of this amino acid in *E. coli*. Finally, the toxin yield was enhanced when phosphopantetheinyl transferase (PPTase) was co-expressed with the *mcy* cluster in *E. coli* (Liu, Mazmouz, and Neilan 2018).

However, MC-LR was not produced directly in this heterologous *E. coli* system due to the lack of the precursor substance D-erythro-β-methyl-iso-aspartic acid. Furthermore, MCs in native systems are originally produced from CO_2_, and studies of both the biosynthetic pathway of MCs and their biological impacts on other microalgal species require autotrophic expression in a model cyanobacterial system. In the current study, we reassembled the *mcy* cluster *in vitro* by TAR cloning, transformed it into the model cyanobacterium *Synechococcus* 7942, and obtained substantial MC-LR production in this model non-toxin-producing cyanobacterium from CO_2_, further functional studies indicate a role in cell division. Our findings lay the foundation for further elucidating the nonribosomal MC biosynthesis pathway in cyanobacteria and suggesting a mechanism of how blooming cyanobacteria gain a competitive edge over their non-blooming counterparts.

## Results

### Assembly of the *mcy* cluster in vitro

A native bidirectional promoter is located between *mcyA* and *mcyD* in the genome of *M. aeruginosa* PCC 7806 (Fig. S1A), which is required for the synthesis of MCs. However, the use of the native bidirectional promoter may be limited in heterologous expression systems, as Liu et al. found that microcystin was not produced using this promoter in *E. coli* (Liu et al. 2017). Here, to successfully produce MC-LR in *Synechococcus* 7942, we used *PpsbA_2_*, a strong light-induced promoter that is commonly used in cyanobacteria, replacing the native bidirectional promoter. Specifically, the bidirectional promoter *biPpsbA_2_* comprises the following (Table S1 and Fig. S2): the flanks of the *biPpsbA_2_* are partial *mcyA* and *mcyD, mcyA’* and *mcyD’* sequences, respectively. The *Sp^r^* resistance gene is located in the middle of *biPpsbA_2_* for the subsequent screening of positive *Synechococcus* 7942 transformants.

We constructed a plasmid containing the *mcy* cluster in yeast strain VL6-48N by TAR cloning (Kouprina and Larionov 2006). We then generated the pGF-NSII-mcy cluster plasmid (75,006 bp) (Fig. S1B) and the control plasmid without the promoter (73,394 bp) (Fig. S1C) via recombination with the genome of *Synechococcus* 7942. The reassembled plasmid maps are shown in Fig. 1.

**Fig. 1.**
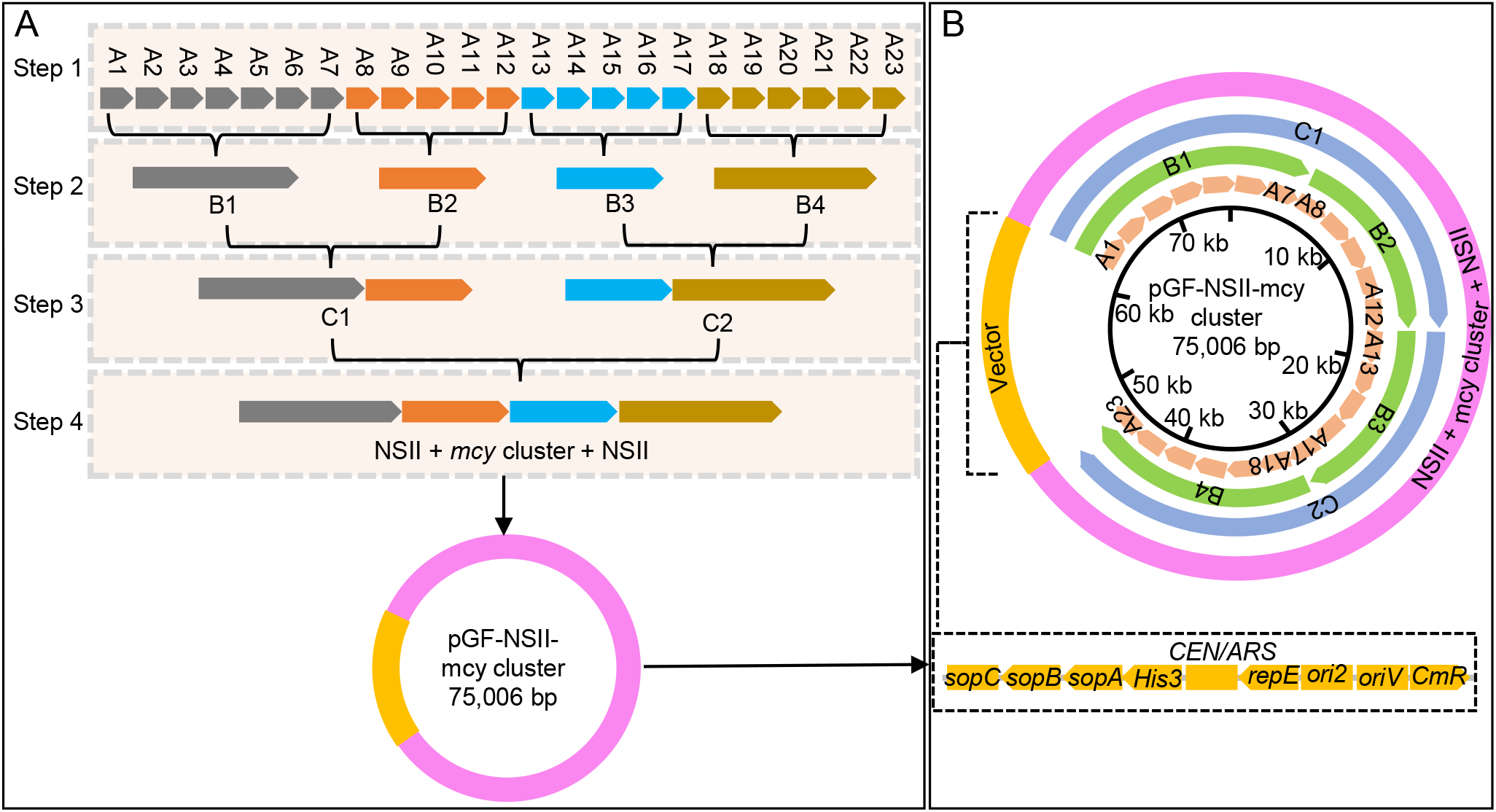
Schematic illustration of the assembly of *mcy* cluster in vitro. (A) Flow diagram of the four steps used to synthesize *mcy* cluster. Each A fragment was ~3 kb, and there was an overlap of no less than 60 bp between two adjacent fragments. Gray, Orange, Blue and Brown arrows represent A1–A7, A8–A12, A13–A17, A18–A23 and their recombination sequence, respectively. (B) A circular map of *mcy* cluster and the corresponding positions of *mcy* cluster and TAR vector region. Light orange arrows represent 23 A fragments in the inner circle; green arrows represent four B fragments in the second layer from the inside; blue arrows represent two C fragments in the third layer from the inside; purple represents *mcy* cluster, *biPpsbA_2_*, and NSII sequence; yellow represents the vector region, sopC, sopB and sopA; His3 is a yeast selectable marker; CEN/ARS are a centromere sequence and origin-of-replication site for plasmid replication in yeast; repE, ori2, and oriV for plasmid replication in *E. coli*; CmR is a selectable marker in *E. coli.*

During the construction of the above plasmid, we performed PCR analysis and Sanger sequencing to verify the sequences of intermediate plasmids B and C and the plasmid pGF-NSII-mcy cluster (primers listed in Table S1). The results indicated that the plasmid was successfully assembled (Fig. 2A–D). Restriction enzyme digestion were further performed to confirm the plasmid sequence. The restriction enzyme digestion profiles of pGF-NSII-mcy cluster were in line with the restriction enzyme profiles simulated using SnapGene 4.1.9 (Fig. 2E and F). High-throughput sequencing also verified the sequence of the plasmid (data not shown). Therefore, the plasmid pGF-NSII-mcy cluster, which contains the *mcy* cluster for the heterologous expression of MC-LR in *Synechococcus* 7942, was successfully assembled in vitro.

**Fig. 2.**
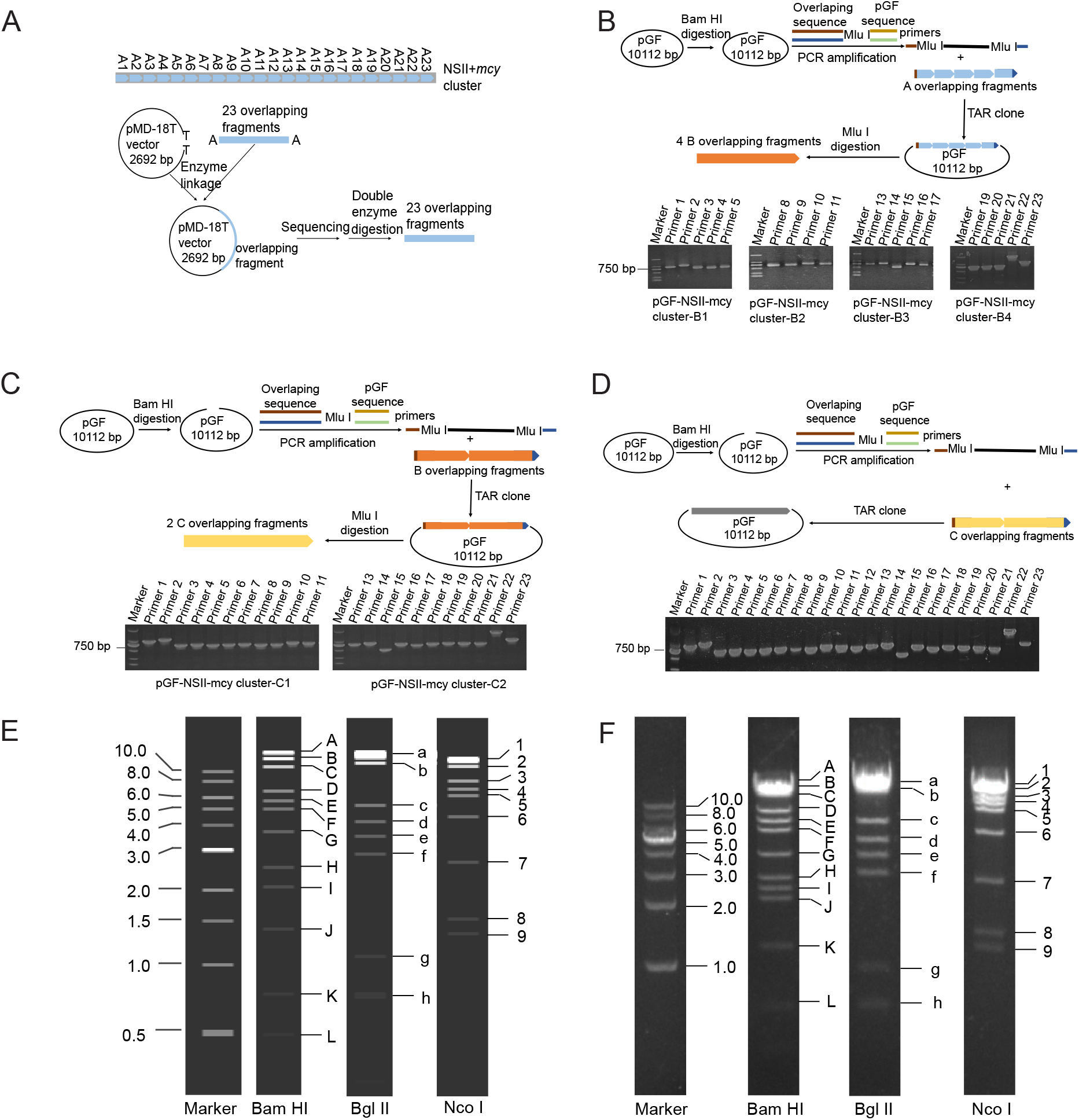
The process and verification used to synthesize the intermediate plasmids and pGF-NSII-mcy cluster. (A) 23 A overlapping fragments (3 kb in size) were acquired by PCR using the genomes of Synechococcus 7942 to NSII, M. aeruginosa PCC 7806 to the *mcy* cluster, and *biPpsbA_2_.* A fragments were used to construct pMD-18T and verified by Sanger sequencing. 23 A fragments were generated by double enzyme digestion. (B) The process used to construct the B plasmids and PCR results. pGF was amplified by PCR, and linearized pGF was used as a template following Bam HI digestion. The primer for pGF vector contained a homologous sequence that was recombined with A fragment using TAR cloning in yeast and an Mlu I site to generate B fragments. (C) The assembly of C plasmids and PCR results. The process is similar to that used for B plasmid construction. (D) pGF-NSII-mcy cluster construction process and PCR results. Gray arrows, *mcy* cluster. Light blue arrows, 23 A fragments. Dark yellow and light green, pGF vector primer sequences. Dark red and dark blue, the overlapping sequences of the A, B, and C fragment on the pGF vector primer. Orange, four B fragments. Yellow, two C fragments. (E) The restriction enzyme profiles of pGF-NSII-mcy cluster predicted by SnapGene 4.1.9 software. (F) The actual restriction enzyme digestion profiles of pGF- NSII-mcy cluster plasmid following 1% agarose gel electrophoresis.

### Heterologous production of MC-LR in *Synechococcus* 7942

We transformed the newly assembled pGF-NSII-mcy cluster into *Synechococcus* 7942 by homologous recombination. The positive transformant, named 7942-NSII-mcy cluster (abbreviation to 7942M), was validated by PCR analysis (Fig. 3A and Table S1) and Sanger sequencing (data not shown). To further verify the transformation of *Synechococcus* 7942 with *mcy* cluster, we performed reverse-transcription PCR (RT-PCR), using *ppc* and *secA* as reference genes. The *mcy* cluster was successfully recombined into the *Synechococcus* 7942 genome and was expressed at the transcriptional level (Fig. 3B and Table S1).

**Fig. 3.**
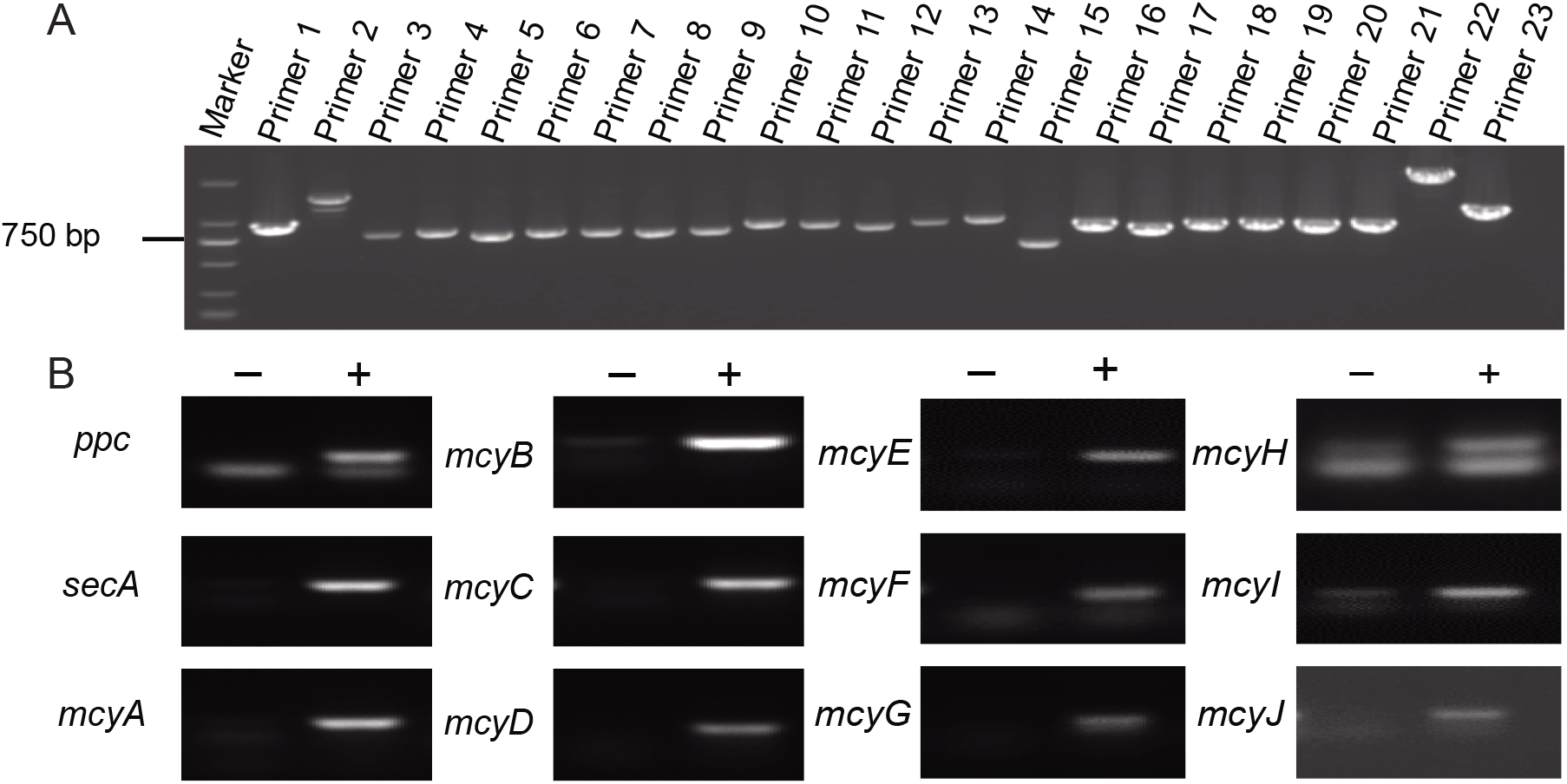
PCR and reverse-transcription PCR (RT-PCR) analysis of the transformant 7942M. (A) PCR analysis; the 7942M genome was used as a template. (B) PT-PCR analysis, cDNA of 7942M was used as a template, represents a sample in which contaminating DNA was removed but the RNA was not reverse transcribed to cDNA, “+” represents PCR results using cDNA.

We cultured 7942M cells and separately extracted intracellular and extracellular microcystin from the cultures. We analyzed the samples by liquid chromatography-mass spectrometry (LC-MS), with the doubly charged [M+2H]^2+^ ion m/z 498 as the parent ion and m/z 135 and m/z 861 as the daughter ions using MC-LR standard and heterologously produced MC-LR (Bruno et al. 2006). The elution time of MC-LR in extracts with m/z 135 and m/z 861 (9.76 min) was consistent with the values in the MC-LR standard at 9.75 min (Fig. 4A–D). We then performed mass spectrometry (MS) to further verify the presence of MC-LR in the extracts. The types of parent ions and main fragmentation ions were consistent between the MC-LR standard and the extracts from the transformant (Fig. 4E–H). Together, these results indicate that the microcystin MC-LR was successfully heterologously produced in the model cyanobacterium *Synechococcus* 7942.

**Fig. 4.**
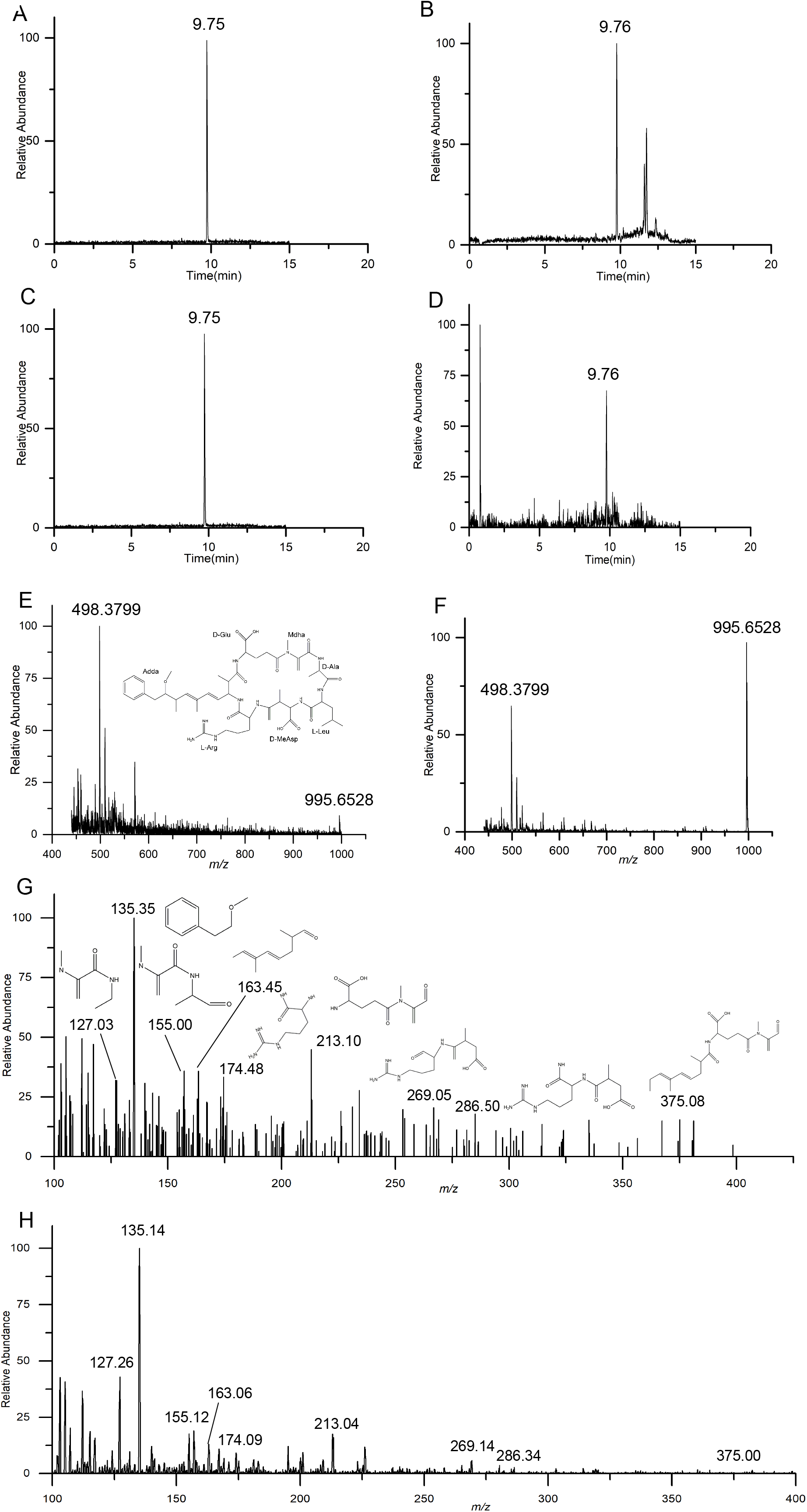
Liquid chromatography-mass spectrometry (LC-MS) and mass spectrometry (MS) analysis of heterologously produced MC-LR from 7942M compared to the MC-LR standard. (A–D) Chromatograms of extracted precursor and fragment ions for (m/z 498.61–135.11) and (m/z 498.61–861.27). (A) MC-LR standard (m/z 498.61–135.11). (B) Heterologously produced MC-LR (m/z 498.61–135.11). (C) MC-LR standard (m/z 498.61-861.27). (D) Heterologously produced MC-LR (m/z 498.61–861.27). (E) and (F) MS spectra from a full-scan of MC-LR standard at 9.75 min (E) and heterologously produced MC-LR at 9.76 min (F), m/z 498 is doubly charged [M+2H]^2+^ ion, m/z 995 is singly charged [M+H] ^+^ ion. (G) and (H) MS spectra of fragmentation ions from m/z 498. (G) MC-LR standard. (H) Heterologously produced MC-LR. The structures shown in (E) and (G) are the structures of precursor and fragment ions.

We confirmed the production efficiency of MC-LR from 7942M by MC-LR-specific (monoclonal primary antibody) ELISA. The yield per gram of dry weight was 6.4 μg g^-1^ for intracellular MC-LR and 0.41 μg L^-1^ for extracellular MC-LR, and the total yield was 10.13 μg g^-1^ (Table 1). The unit cell productivity from 7942M was 0.006–0.018 fg cell^-1^ day^-1^.

**Table 1.** Yield of MC-LR from 7942M, as determined by MC-LR-specific ELISA

| MC-LR | Yield |
| --- | --- |
| Intracellular ($\mu\text{g g}^{-1}$) | 6.4±0.007 |
| Extracellular ($\mu\text{g L}^{-1}$) | 0.41±0.02 |
| Total ($\mu\text{g g}^{-1}$) | 10.13±0.02 |

All data points in the table represent the means of three replicated ELISAs of each independent culture, with the SD of the means $P<0.05$ or $P<0.01$ ).

MCs irreversibly inhibit type 1 and 2A eukaryotic serine/threonine protein phosphatases (PP1/PP2A) (Yoshizawa et al. 1990; Mackintosh et al. 1990), causing an imbalance in cell phosphorylation levels and metabolic disorders in the cell (Falconer and Humpage 2005; Sotton et al. 2012). We therefore performed a PP2A assay to confirm the biological activity of heterologously expressed MC-LR. The PP2A inhibitory activities of MC-LR standard and MC-LR from 7942M were similar (Fig. 5). Therefore, the heterologously produced MC-LR maintained the same biological activity as natural MC-LR from *M. aeruginosa* PCC 7806.

**Fig. 5.**
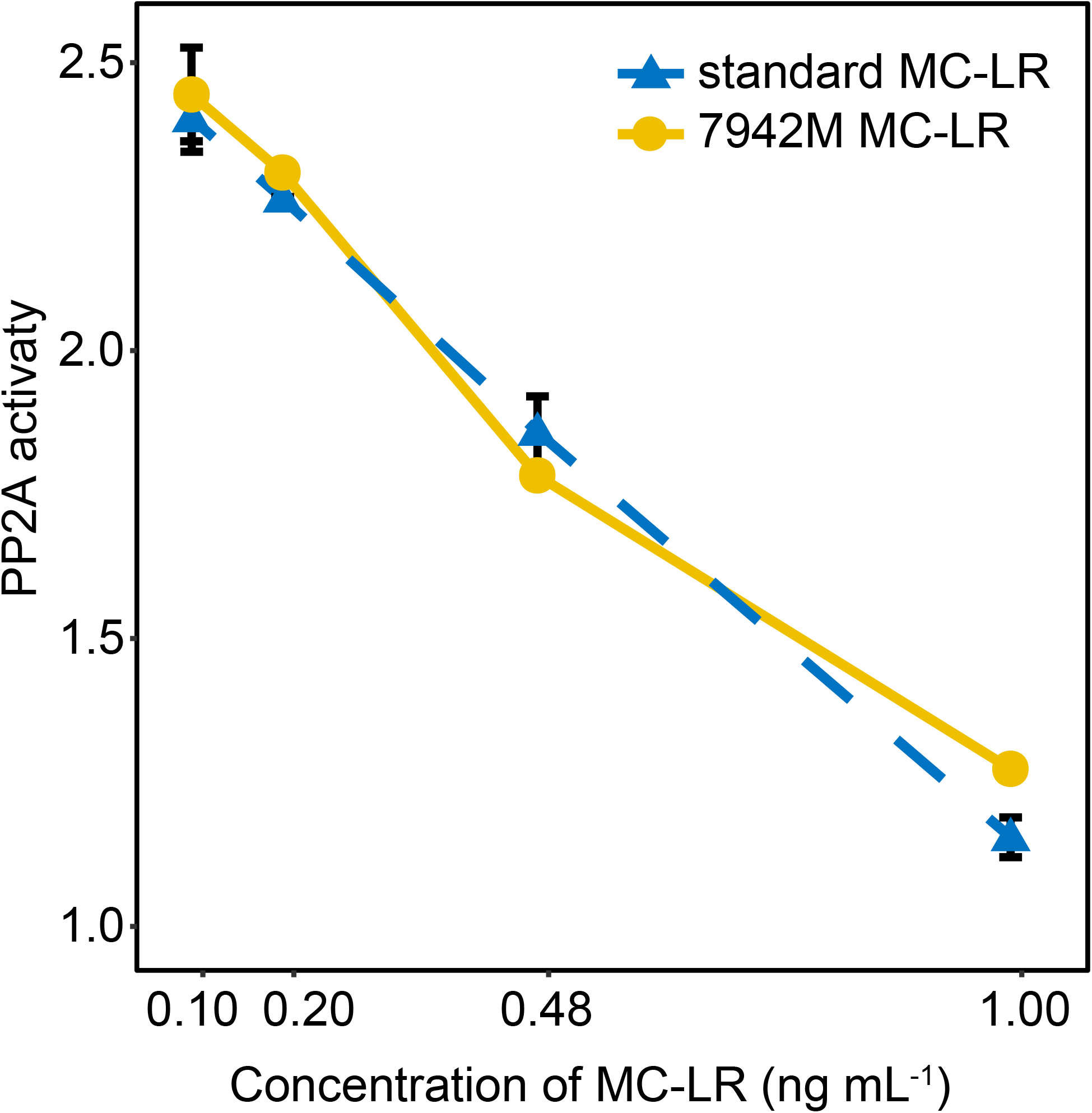
Comparison curve of the inhibitory activity of protein phosphatase 2A (PP2A) between MC-LR standard and heterologously expressed MC-LR from a 7942M extract.

### Heterologous production of MC-LR inhibits *Synechococcus* 7942 cell division

Cell morphology observed by fluorescence microscopy shows that as compared with the WT and the control, the 7942M cells are obviously filamentous with polarized photosynthetic pigments distribution (indicated by arrows in Fig. 6A and Fig. S4) and shows abnormal cell division throughout the culturing period (Fig. 6A and Fig. S3A), and the WT cells show similar phenotype to that of the 7942M when artificially fed with MC-LR (Fig. 6A and Fig. S3B). A total count of 1664 WT cells and 1925 7942M cells were chosen to calculate the cell size distribution, the length of WT cells varies between 1.42 and 5.95 μm with 3.2 μm at the most (the longer ones are mostly in division), while that of the 7942M ranged from 1.04 to 58.9 μm with 8.07 μm accounted for the most (Fig. 6B).

**Fig. 6.**
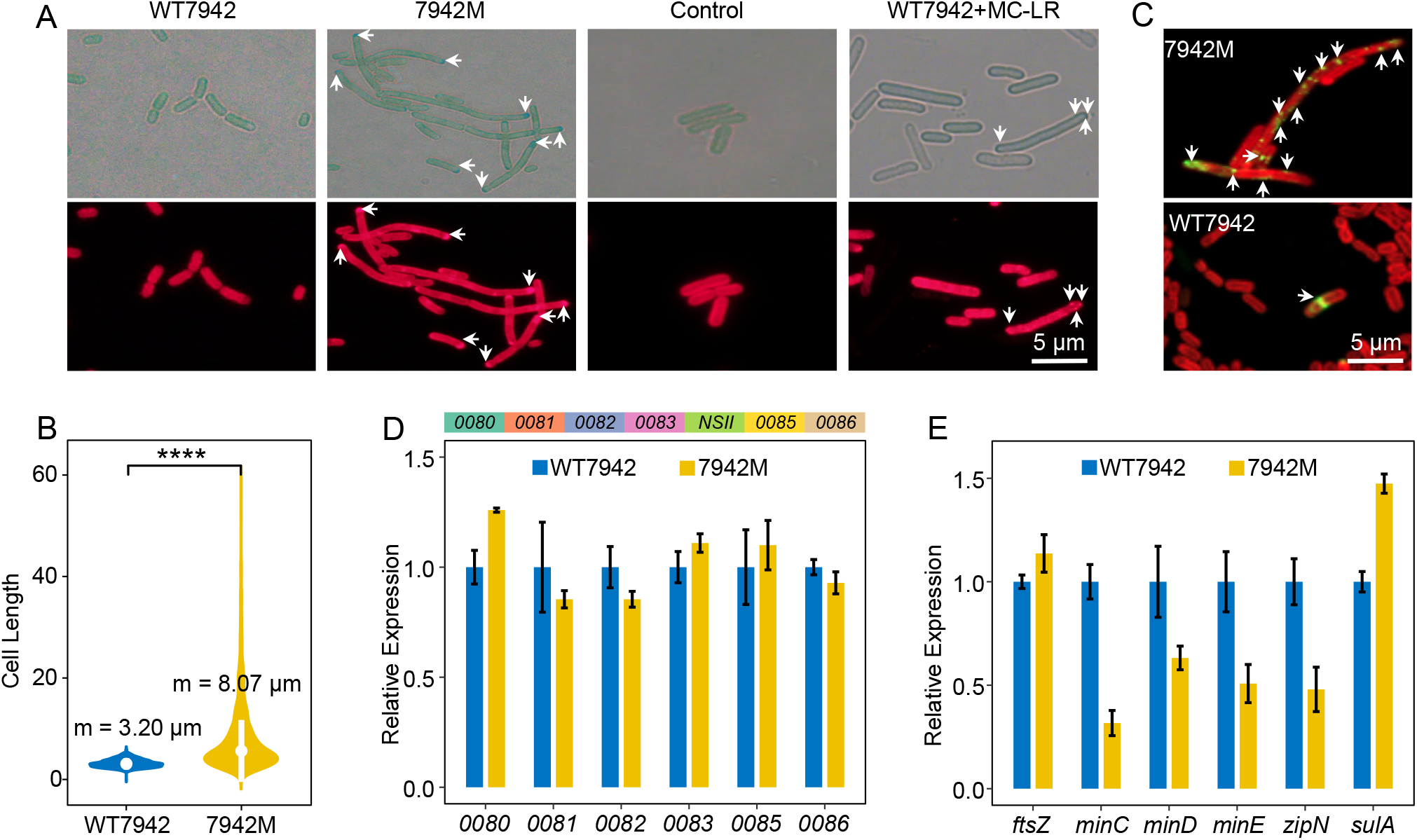
Cell morphology changes, FtsZ localization and the expression of related genes. (A) Microscopic images of cell morphology changes in WT7942, 7942M, control and WT7942+MC-LR shown in bright field (upper images), and auto-fluorescence (lower images), arrows in those 7942M and WT7942+MC-LR images showing aggregated pigments. (B) The distribution of cell sizes in WT7942 and 7942M. (C) Immunofluorescence analysis showing localization of FtsZ in 7942M (upper images) and WT7942 (lower images). (D) Distribution map of genes around *NSII* (upper) and expression level of gene around *NSII* (lower). (E) Expression level of gene related to cell division.

The FtsZ protein is a tubulin-like GTPase protein which would assemble Z-ring structure as the skeleton and scaffold to the cellular divisome (Zhang et al. 2020; Dong et al. 2010). The localization of FtsZ were examined by immunofluorescence microscopy, as shown in Fig. 6C and Fig. S4, the Z-ring was correctly localized in midcell in WT cells as expected, which were hardly seen in 7942M and FtsZ proteins seems to be aggregated.

In order to study whether the cell filamentation of 7942M is caused by the recombination of the *mcy* cluster on the genome of *Synechococcus* 7942 or by malfunctioned cell division, we detected the expression level of the related genes, including *synpcc7942_0080* (*0080*), *synpcc7942_0081* (*0081*), *synpcc7942_0082* (*0082*), *synpcc7942_0083* (*0083*), *synpcc7942_0085* (*0085*), *synpcc7942_0086* (*0086*) surrounding the *NSII* site, and those genes encoding proteins responsible for cell division, such as FtsZ, Min system, including MinC, MinD and MinE, which regulate the assembly of Z-ring (Jordan et al. 2017); ZipN, the essential component for divisome assembly by interactings with FtsZ and other central divisome components (Camargo et al. 2019); SulA, regulates the assembly and position of Z-ring structure by directly interacting with FtsZ (Koksharova and Babykin 2011). While as expected, the expression levels of those genes surrounding *NSII* are unaffected (Fig. 6D and Table S1). Although the gene expression level of *ftsZ* in 7942M is similar to that of the WT, the expression levels of the *min* system and *zipN* are significantly down-regulated, with *sulA* being up-regulated in 7942M (Fig. 6E and Table S1). The results indicated that the production of MC-LR in *Synechococcus* 7942 leads to malfunctioned cell division, by affecting the functional assembly and/or positioning of the cell division protein FtsZ.

### MC-LR competes the GTP binding sties of FtsZ

To further distinguish the changes of the expression level of the cell division related genes are whether causes or results of the cell filamentation in 7942M, and to understand if MC-LR directly interacts with FtsZ and how, we predicted the crystal structure of *Synechococcus* 7942 FtsZ using I-TASSER by referencing all published FtsZ structures, and carried out molecular docking analysis to predict possible interactions (Vakser 2014) using Auto dock vina, Auto Dock tools1.5.6, Pymol v.2.3 and UCSF Chimera1.14. Sequence alignment (Fig. S5) shows that like other FtsZs, *Synechococcus* 7942 FtsZ has two conserved elements, GG*N (47-51) and GGTG (133-136), which form the nucleic acid binding domain (NBD) together with the conserved THR^159^ and ARG^169^. As shown in Fig. S6, both GTP (Fig. S6A) and MC-LR (Fig. S6B) could be fitted into the same binding pocket of FtsZ monomer (Fig. S6C and D), and compete binding amino acid residues GLY^47^, GLY^48^, and THR^159^ (Fig. S6E and F), with GLY^47^, GLY^48^ located within the conserved GG*N (47-51) NBD (Fig. S5). FtsZ starts polymerization when combined with GTP (Olson, Wang, and Osteryoung 2010). Fig. S7 shows that both GTP (Fig. S7A) and MC-LR (Fig. S7B) could also be localized into the same binding pocket of FtsZ dimer (Fig. S7C and D), compete binding the conserved amino acid residues GLY^47^, GLY^48^, THR^159^ and ARG^169^ (Fig. S7E and F). The increased docking score and the changed interaction amino acid residues in dimeric FtsZ (Fig. S6 and S7) indicate a conformational change of the binding pocket, in accordance with the high affinity FtsZ conformation and low affinity FtsZ conformation observed by Miraldi et al (Miraldi, Thomas, and Romberg 2008).

MicroScale Thermophoresis (MST) is an immobilization-free technology for the biophysical analysis of interactions between biomolecules. In order to verify the molecular docking predictions, the interactions between GTP and FtsZ, MC-LR and FtsZ, the competitive binding of GTP and MC-LR to FtsZ, were analyzed respectively through MST. Fig. 7 shows that the MST analyzed Kd value of GTP and MC-LR for FtsZ are 15.2±8.6 nM (Fig. 7A) and 1.12±0.65 μM (Fig. 7B), respectively. However, when FtsZ was first combined with MC-LR, the binding affinity of GTP to FtsZ decreased by 23 times to 347±152 nM (Fig. 7C). FtsZ polymerization were further investigated by 90° angle light scattering (Mukherjee and Lutkenhaus 1999), in the presence of GTP, MC-LR, and both, the results again proved that the binding of MC-LR to FtsZ greatly reduces the FtsZ polymerization triggered by GTP binding (Fig. 7D). These results proved that MC-LR does irreversibly compete for the GTP binding site of FtsZ, thereby reduces the rate of FtsZ polymerization, makes the daughter cells unable to separate normally, and results in cell filamentation (Fig. 6A), which proved that the abnormal cell division related gene expression (Fig. 6E) are just results, not causes.

**Fig. 7.**
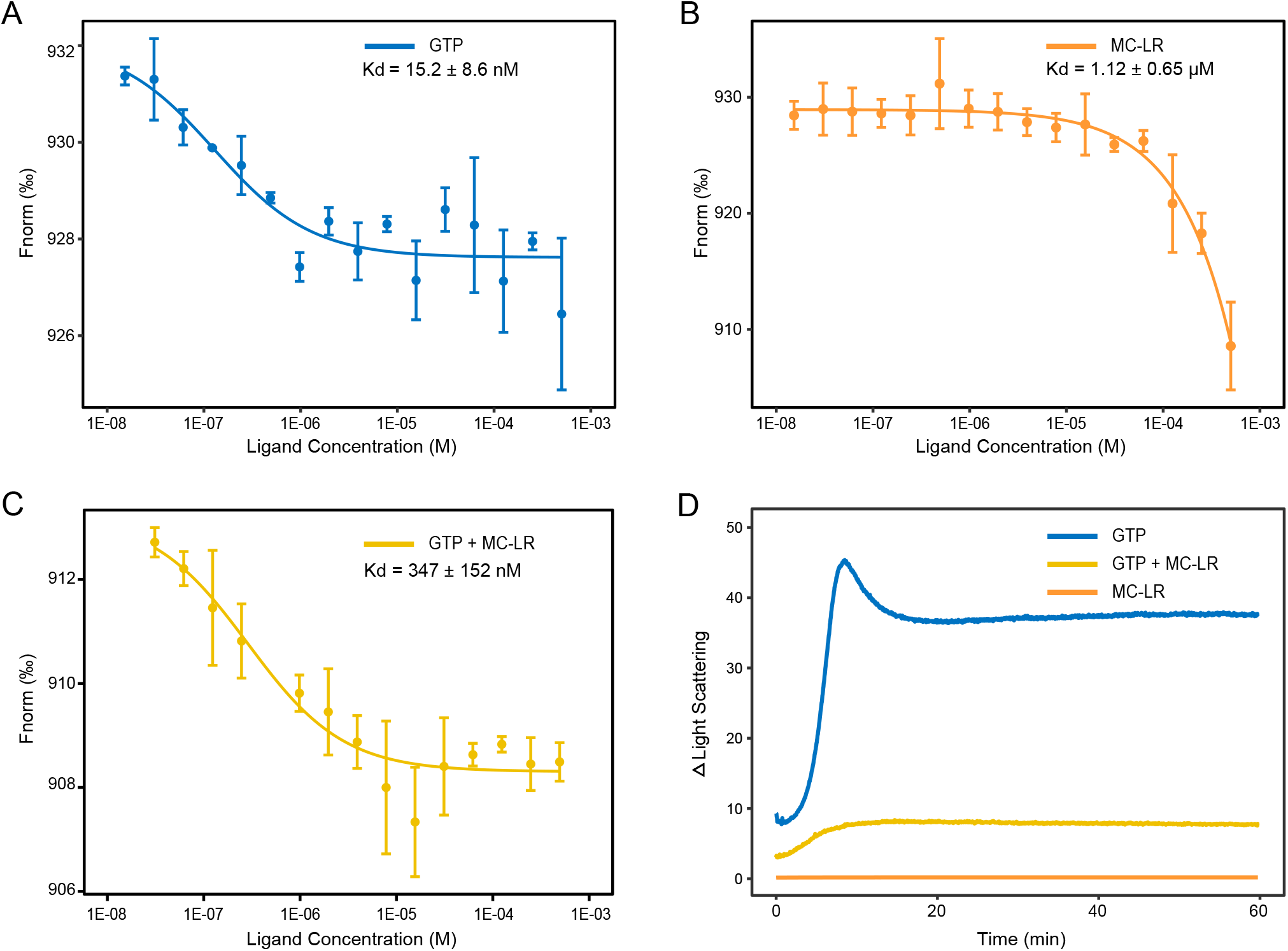
MST and 90° angle light scattering analysis of the interaction of FtsZ with GTP and MC-LR. (A) FtsZ-GTP interaction (Kd = 15.2 ± 8.6 nM). (B) FtsZ-MC-LR interaction (Kd = 1.12 ± 0.65 μM). (C) Competitive MST, FtsZ was first labeled by fluorochrome and incubated briefly with MC-LR, the affinity of GTP to FtsZ reduced to 347 ± 152 nM. (D) FtsZ polymerization investigated by 90° angle light scattering, in the presence of GTP (blue), MC-LR (orange), and both (yellow).

## Discussion

Assembling large fragments of genomic DNA in vitro remains a technical challenge for synthetic biology. The three most common methods currently used for this process are artificial synthesis, fosmid library construction, and TAR cloning, which have successfully been used to assemble 770 kb, 35.6 kb, and 300 kb fragments, respectively (Kouprina and Larionov 2006; Jiang et al. 2020; Liu et al. 2017). Direct artificial synthesis in vitro can be used for codon optimization. For example, the yeast chromosome (Sc2.0 Project) (12 Mb) had been over half synthesized in vitro using artificial synthesis technology (Jiang et al. 2020). However, the high cost of this technique limits its application. Fosmid library construction has been successfully used for the heterogeneous expression of microcystin in *E. coli* (Liu et al. 2017). However, the process of genomic fosmid library construction is difficult to control, labor-intensive, and requires a large number of repetitive homologous recombination processes for fragment connection. By contrast, the TAR cloning technique is simple, requires little time, leads to high rates of recombination, and can be used to synthesize DNA sequences of up to 300 kb (Noskov et al. 2001). Shang et al. used this technique to synthesize the 145,299 bp baculovirus genome in vitro and used it to transfect Sf9 insect cells (Shang et al. 2017).

Liu and co-workers synthesized two microcystin variants (MC-LR and [D-Asp^3^]- MC-LR) in *E. coli* (Liu et al. 2017). However, since D-erythro-β-methyl-iso-aspartic acid, the precursor substance of position 3 of MCs, is absent in *E. coli*, MC-LR was not produced directly. In addition, Red/ET recombineering was used to connect two libraries, but this technique is difficult to perform in cyanobacteria. In the present study, we synthesized heterogeneous MC-LR in a photosynthetic model cyanobacteria without adding any substrate. After successfully assembling the pGF-NSII-mcy cluster (Fig. 1 and 2) in vitro by TAR cloning, the *mcy* cluster was transformed into *Synechococcus* 7942 (Fig. 3), and heterologously expressed MC was synthesized (Fig. 4 and 5). In addition, we assembled the autonomously replicating plasmid pGF-mcy cluster (73,818 bp) (Fig. S1D, S8 and S9) and transformed it into *Synechococcus* 7942 by conjugational transfer (Chen, van der Steen, et al. 2016). The positive transformant, named 7942-mcy cluster, was validated in the first generation (Fig. S10A). However, the autonomously replicating plasmid pGF-mcy cluster was degraded in the fifth generation transformant (Fig. S10B), and plasmid degradation also occurred in the reactivated first-generation transformant following transfer from an ultra-low temperature freezer (Fig. S10C).

RSF1010-derived plasmids are most efficiently transferred into *Synechocystis* 6803 and 6714 and *Synechococcus* 7942 and 6301 by conjugation, which can even replicate automatically without cyanobacterial DNA (Mermet-Bouvier et al. 1993; Zinchenko et al. 1999; Savakis, Angermayr, and Hellingwerf 2013; Chen, Taton, et al. 2016). However, although the self-replicating plasmid pGF-mcy cluster containing replication-related genes from RSF1010 replicated and survived in *Synechococcus* 7942, it was genetically unstable (Fig. S10 B and C). Two hypotheses could explain this instability: expression pressure and replication pressure of large DNA fragments. Specifically, it is challenging to express the toxic *mcy* cluster in this self-replicating plasmid due to expression pressure. Alternatively, the presence of replication-related genes from RSF1010 may not have been sufficient to support the replication of the large (73,818 bp) pGF-mcy cluster plasmid, as it is difficult to replicate such large plasmids. Therefore, compared to self-replicating plasmids, the recombination of the *mcy* cluster into the genome of *Synechococcus* 7942 is more suitable for the heterologous production of microcystin.

The yield of MC-LR from *M. aeruginosa* PCC 7806 was 0.65–2.2 fg cell^-1^ day^-1^ (Jahnichen et al. 2007), and the yield of MC-LR in the Fosmid-*E. coli* produced via the synthesis of the *mcy* cluster was 0.0007–0.0059 fg cell^-1^ day^-1^ (Liu et al. 2017). In the current study, MCs were expressed in the model non-toxic cyanobacterium *Synechococcus* 7942 via TAR cloning and assembly in vitro, yielding recombinant MC-LR at a rate of 0.006–0.018 fg cell^-1^ day^-1^. And the biological activity of MC-LR reassembled in 7942M was confirmed by PP2A assay (Fig. 5). Although the unit cell yield of 7942M was lower than that of *M. aeruginosa* PCC 7806, it was one order of magnitude higher than that of the Fosmid-*E. coli*. Furthermore, compared with *M. aeruginosa* PCC 7806, it is easier to carry out genetic manipulation in model cyanobacteria, making it convenient to study the biosynthetic process of MC-LR and its role in cyanobacterial blooms. Compared with heterologous expression in *E. coli* (Liu et al. 2017), this process is closer to the biosynthetic process of this compound in *M. aeruginosa* PCC 7806. Importantly, in the current study, we performed the first synthetic biological expression of the *mcy* cluster and demonstrated the autotrophic production of MC-LR in a photosynthetic model organism.

The MC-LR producing mutant strain 7942M fails to assemble the Z-ring and shows filamentous cell (Fig. 6A–C) with unaffected FtsZ expression (Fig. 6E), molecular docking analysis of GTP and MC-LR to FstZ indicated that MC-LR could be perfectly docked into the GTP binding domain of both FtsZ monomer and dimer (Fig. S6 and S7), and better as dimer, although reacted with different linker amino acid residues. The differences of the docking affinity and binding pocket conformational changes between the FtsZ monomer and the dimer, are in accordance with the observed low and high affinity conformation dominated by FtsZ monomer and polymer(Miraldi, Thomas, and Romberg 2008), and partially proved the hypothesis of the shift or rotation of the two subdomains(Chen and Erickson 2011) when FtsZ first polymerized to dimers(Chen et al. 2005). Further MST and 90° angle light scattering experiments proved that MC-LR irreversibly competes with the GTP binding domain of FtsZ and prevents it from polymerization (Fig. 7). The results indicated that by competing with the GTP binding site in FtsZ, MC-LR inhibits the formation of FtsZ protofilaments, and resulted in undivided long filamentous cells in 7942M, which could possibly be one good reason that blooming cyanobacteria gain a competitive edge over their non-blooming counterparts.

In conclusion, We reassembled the pGF-NSII-mcy cluster in vitro by TAR cloning and recombined *mcy* cluster into the *NSII* site of the *Synechococcus* 7942 genome to obtain bioactive MC-LR, and proposed the following working model for the competition between GTP and MC-LR to the binding site of FtsZ (Fig. 8): Normally, the binding of GTP to FtsZ monomer initiate the polymerization of FtsZ to its dimeric high affinity conformation, the later will then further cooperatively assembled into protofilament and finally leads to cell division; however, in the presence of MC-LR, which irreversibly competing the GTP binding pocket of FtsZ, both monomer and dimer, hold it back from further polymerization and thus results in abnormal cell division in *Synechococcus* 7942.

**Fig. 8.**
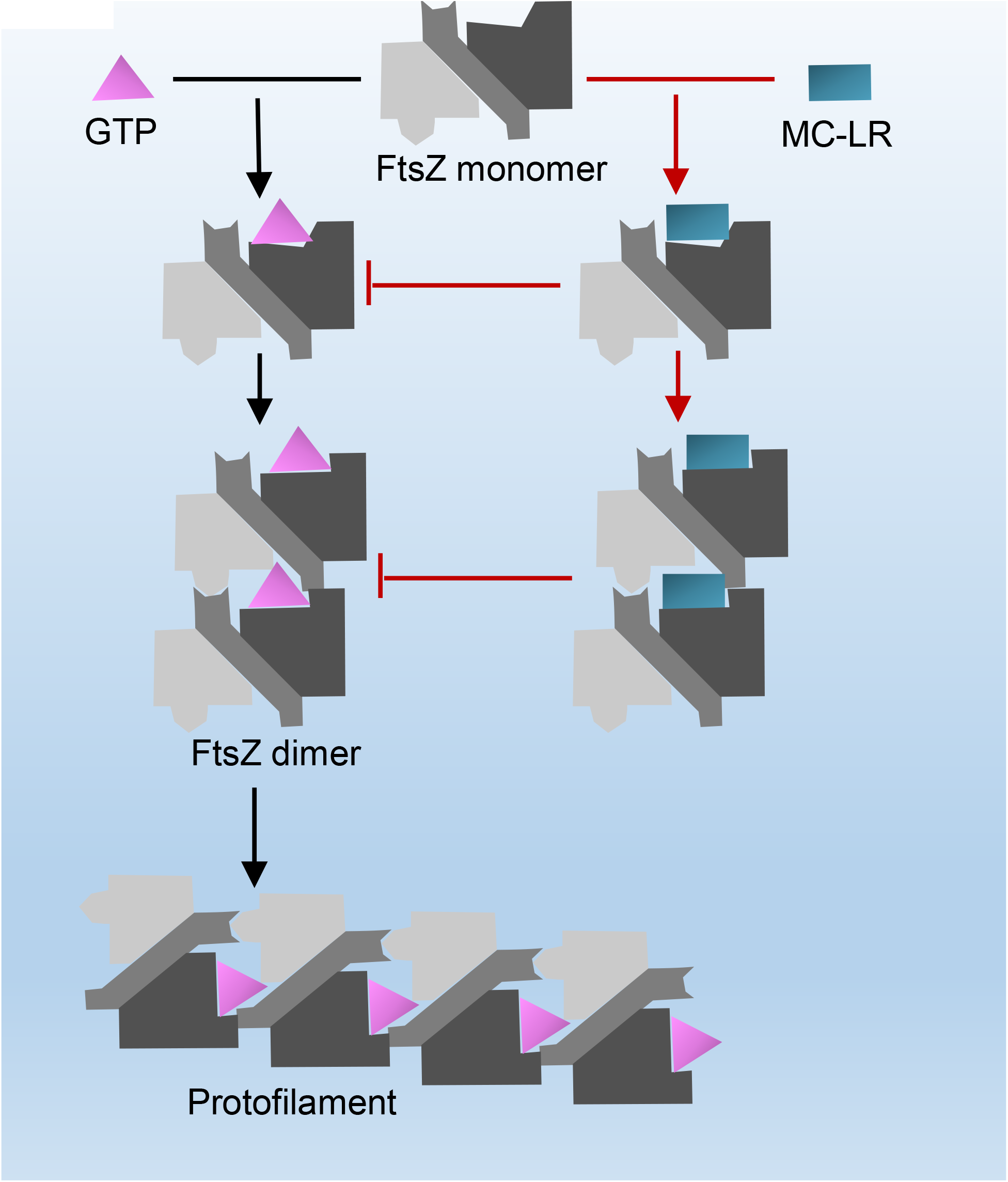
Proposed model for the competition between GTP and MC-LR to the NBD site of FtsZ. The binding of GTP to FtsZ monomer (low affinity conformation) initiate the polymerization of FtsZ to its dimeric high affinity conformation, the later will then further cooperatively assembled into protofilament and finally leads to cell division; however, in the presence of MC-LR, which irreversibly competing the GTP binding pocket of FtsZ, both monomer and dimer, hold it back from further polymerization and thus results in abnormal cell division in 7942M.

## Materials and Methods

### Yeast strain, E. coli strain, algae strains, and culture conditions

*S. cerevisiae* strain VL6-48 (*MAT alpha, his3-Δ200, trp1-Δ1, ura3-Δ1, lys2, ade2-101, met14), E. coli* strain EPI300 carrying an inducible *trfA* gene, and the TAR cloning vector pGF were obtained from the Research Group of Systems Virology, State Key Laboratory of Virology of the Wuhan Institute of Virology, Chinese Academy of Sciences (Hou et al. 2016; Shang et al. 2017). *M. aeruginosa* PCC 7806 was purchased from the Freshwater Algae Culture Collection at the Institute of Hydrobiology (FACHB) and cultured in BG11 medium with constant illumination at 60 μmol photons m^-2^ s^-1^ at 30°C. Wild-type *Synechococcus* 7942 (WT7942) (from our laboratory), 7942-NSII-mcy cluster (7942M), control and 7942-mcy cluster (generated in this study) were cultured in BG11 medium with constant illumination at 80 μmol photons m^-2^ s^-1^ at 30°C. The cells were observed using a fluorescence microscope (Olympus BX53, Japan).

### Construction of the bidirectional promoter *biPpsbA_2_*

The *biPpsbA_2_* promoter was constructed by PCR amplification and restriction enzyme digestion (Fig. S2). *mcyD* and *mcyA’* (the homologous sequences of *mcyD* and *mcyA*, respectively) were recombined with the light-inducible strong promoter *PpsbA_2_* by PCR fusion. The recombination fragments were inserted into the linearized pMD-18T vector (verified by Sanger sequencing), generating the plasmids D-PpsbA_2_ (pD-PpsbA_2_) and A-PpsbA_2_ (pA-PpsbA_2_). The Spectinomycin resistance gene (*Sp^r^*) fragment was generated by digesting the pRL57 plasmid with Dra I. The *Sp^r^* gene fragment was inserted into the linearized pD-PpsbA_2_ plasmid by Sal I digestion to generate D-PpsbA_2_-Sp^r^ (pD-PpsbA_2_-Sp^r^). pD-PpsbA_2_-Sp^r^ was double digested with Kpn I and Sph I to generate the D-PpsbA_2_-Sp^r^ fragment, which was introduced into linearized pA-PpsbA_2_ by Kpn I digestion, generating pD-PpsbA_2_-Sp^r^-PpsbA_2_-A. Finally, pD-PpsbA_2_-Sp^r^-PpsbA_2_-A was digested with Sac I and Pst I to produce the bidirectional promoter *biPpsbA_2_*. All plasmids were confirmed by PCR amplification, and the primers are shown in Table S1.

### Synthesis and construction of the pGF-NSII-mcy cluster, the control and the pGF-mcy cluster plasmids

The TAR cloning vector pGF was used to construct the pGF-NSII-mcy cluster (Fig. S1B), the control (Fig. S1C) and the pGF-mcy cluster (Fig. S1D) plasmids (Hou et al. 2016; Kouprina and Larionov 2008; Shang et al. 2017). The pGF-NSII-mcy cluster was synthesized based on TAR in yeast via four steps, as outlined in Fig. 1. The detailed recombination process to generate the pGF-NSII-mcy cluster plasmids is summarized in Fig. 2A–D.

First, the *mcy* cluster from *M. aeruginosa* PCC 7806, the bidirectional promoter *biPpsbA_2_* (A15), and the neutral site (NSII) gene from *Synechococcus* 7942(Zhan et al. 2016) were amplified into 23 overlapping fragments (A1-A23) (Table S1) and introduced into the pMD-18T vector. Following confirmation by Sanger sequencing, 23 overlapping fragments were generated by double enzyme digestion (Fig. 2A).

Second (Fig. 2B), the pGF vector was linearized by Bam HI digestion. Two overlapping sequences were added to two long primers, pGF-A7-F and pGF-A1-R: the 77 nucleotide (nt) pGF-A7-F sequence contained A7 followed by a Mlu I site and 17 nt of pGF downstream of the Bam HI site; the 74 nt pGF-A1-R sequence contained A1 followed by a Mlu I site and 20 nt of pGF upstream of the Bam HI site. The pGF vector was amplified by PCR using two pairs of primers (pGF-F and pGF-A1-R, pGF-R and pGF-A7-F) to generate two vector fragments. Yeast protoplasts were cotransformed with the two vector fragments and fragments A1–A7, generating the intermediate plasmid pGF-NSII-mcy cluster-B1. Yeast protoplasts were then cotransformed with two vector fragments carrying the 46 nt A12 sequence and the 48 nt A8 sequence and fragments A8–A12, generating the intermediate plasmid pGF-NSII-mcy cluster-B2. Yeast protoplasts were cotransformed with two vector fragments carrying the 46 nt sequence A17 and the 48 nt sequence A13 and fragments A13–A17, generating the intermediate plasmid pGF-NSII-mcy cluster-B3. Yeast protoplasts were then cotransformed with two vector fragments carrying the 77 nt sequence A23 and the 48 nt sequence A18 and fragments A18–A23, generating the intermediate plasmid pGF-NSII-mcy cluster-B4. The four intermediate B plasmids were extracted from the yeast cells and transformed into *E. coli* EPI300 for further amplification. The four amplified B plasmids were digested with Mlu I to generate four B fragments.

Third (Fig. 2C), the intermediate plasmids pGF-NSII-mcy cluster-C1 and pGF-NSII-mcy cluster-C2 were generated in a similar manner to pGF-NSII-mcy cluster-B. Yeast protoplasts were cotransformed with two vector fragments carrying the 46 nt A12 sequence and the 74 nt A1 and B1 sequences with the B2 fragments, generating the intermediate plasmid pGF-NSII-mcy cluster-C1. Yeast protoplasts were cotransformed with two vector fragments carrying the 77 nt A23 sequence and the 48 nt A13 and B3 sequences with the B4 fragments, generating the intermediate plasmid pGF-NSII-mcy cluster-C2.

Finally (Fig. 2D), yeast protoplasts were cotransformed with two vector fragments carrying the 77 nt A23 sequence and the 74 nt A1 and C1 sequences with the C2 fragments, generating the plasmid pGF-NSII-mcy cluster, with a total length of 75,006 bp. The primers used for pGF amplification are shown in Table S1.

The vector fragments and intermediate fragments used for TAR cloning were quantified throughout the cloning process. In steps 2 and 3, all intermediate plasmids were confirmed by PCR amplification (Table S1) and Sanger sequencing (data not shown). In step 4, the plasmid pGF-NSII-mcy cluster was identified by PCR amplification (Table S1), Sanger sequencing (data not shown), restriction enzyme (Bam HI, Bgl II, and Nco I) analysis (Fig. 2E and F), and high-throughput sequencing (by TSINGKE Biological Technology Co., Ltd., China) (data not shown).

The control plasmid was constructed following the same protocol as the pGF-NSII-mcy cluster, except that there are no promotors between *mcyA* and *mcyD*.

The pGF-mcy cluster was synthesized in a similar manner to the pGF-NSII-mcy cluster. The process is shown in Fig. S8. The difference is that 23 A’ fragment of pGF-mcy cluster (Table S1) included the *mcy* cluster from *M. aeruginosa* PCC 7806, the bidirectional promoter *biPpsbA_2_*, and replication-related genes from RSF1010. Yeast protoplasts were cotransformed with two vector fragments carrying the 77 nt A5’ sequence (equivalent to A7 of pGF-NSII-mcy cluster), the 74 nt A1’ sequence (Table S1) and the A1’–A5’ fragments, generating the intermediate plasmid B1’. Yeast protoplasts were then cotransformed with vector fragments carrying the 46 nt A10’ sequence (equivalent to A12 of pGF-NSII-mcy cluster), the 48 nt A6’ sequence (equivalent to A8 of pGF-NSII-mcy cluster), and the A6’–A10’ fragments, generating the intermediate plasmid B2’. Yeast protoplasts were cotransformed with two vector fragments carrying the 46 nt A15’ sequence (equivalent to A17 of pGF-NSII-mcy cluster), the 48 nt A11’ sequence (equivalent to A13 of pGF-NSII-mcy cluster), and the A11’–A15’ fragments, generating the intermediate plasmid B3’. Yeast protoplasts were then cotransformed with two vector fragments carrying the 46 nt A20’ sequence (Table S1), the 48 nt A16’ sequence (equivalent to A18 of pGF-NSII-mcy cluster), and the A16’–A20’ fragments, generating the intermediate plasmid B4’. Finally, yeast protoplasts were cotransformed with two vector fragments carrying the 77 nt A23’ sequence, the 73 nt A21’ sequence, and the A21’–A23’ fragments, generating the intermediate plasmid B5’.

Second, yeast protoplasts were cotransformed with two vector fragments carrying the 46 nt A15’ sequence, the 74 nt A1’ sequence, and the B1’–B3’ fragments, generating the intermediate plasmid C1’. Yeast protoplasts were cotransformed with two vector fragments carrying the 77 nt A23’ sequence, the 48 nt A16’, B4’ sequences and B5’ fragments, generating the intermediate plasmid C2’.

Finally, yeast protoplasts were cotransformed with two vector fragments carrying the 77 nt A23’ sequence, the 74 nt A1’ and C1’ sequences, and the C2’ fragments, generating the intermediate plasmid pGF-mcy cluster, with a total length of 73,818 bp.

All intermediate plasmids were confirmed by PCR amplification (Fig. S9A upper and middle) and Sanger sequencing (data not shown). The plasmid pGF-mcy cluster was identified by PCR amplification (Fig. S9A lower), Sanger sequencing (data not shown), restriction enzyme (Bam HI and Nde I) (Fig. S9B) analysis, and high-throughput sequencing (by TSINGKE Biological Technology Co., Ltd., China) (data not shown).

### Transformation of *Synechococcus* 7942 with pGF-NSII mcy-cluster and pGF-mcy cluster

The pGF-NSII-mcy cluster plasmid was transferred into *Synechococcus* 7942 cells by homologous recombination(Ma et al. 2017). A 10–mL aliquot of cells (OD_730_ < 1.0) was washed three times with 10 mM NaCl, centrifuged at 2,000 g for 10 min at room temperature, and resuspended in 200 μL fresh BG11 medium. The cells were combined with 20 μL of 100 ng μL^-1^ pGF-NSII-mcy cluster and cultured under low-light conditions for 8-10 h. A 100 μL aliquot of culture was plated onto a 0.2 μm nitrocellulose membrane on a non-resistant BG11 plate. After 20–24 h of incubation at 30°C, the nitrocellulose membrane was transferred to a BG11 plate containing Spectinomycin.

The pGF-mcy cluster was transferred to *Synechococcus* 7942 via tri-parental mating(Chen, van der Steen, et al. 2016). *E. coli* RP4 (from Research Group of Algal Genetics and Biotechnology, Institute of Hydrobiology, Chinese Academy of Sciences) was used as a helper strain, and *E. coli* EPI300 harboring pGF-mcy cluster was used as a donor strain. For each strain, 10 mL LB without antibiotics was inoculated with 250 μL of an overnight culture. After 2.5–3 h of incubation, the cultures were harvested by centrifugation at 800 g for 8 min at room temperature. The pellets of the donor and helper strains were each resuspended in 1–mL fresh LB, mixed, and concentrated via centrifugation into 100 μL fresh LB. The mixture was incubated at 30°C for 1 h, followed by the addition of 800 μL of a young culture (OD_730_ < 1.0) of *Synechococcus* 7942. The three-strain mixture was centrifuged, 800 g, 8 min at room temperature and the pellet resuspended in 30 μL fresh BG11 medium. The cells were spread onto the surface of a 0.2 μm nitrocellulose membrane on a non-resistant BG11 plate with 5% (v/v) LB medium. Following incubation under low-light conditions for 20–24 h at 30°C, the nitrocellulose membrane was transferred to a BG11 plate containing Spectinomycin and incubated at 30°C under normal illumination.

### Reverse Transcription PCR analysis

RNA was extracted from WT7942 harboring 7942M as described by Hu et al.(Hu et al. 2018). cDNA was synthesized using a HiScript II Q RT SuperMix for qPCR (+gDNA wiper) kit (# R223-0, Vazyme, China), and 4× gDNA Wiper Mix and 5× No RT Control Mix from this kit were used to remove contaminating DNA; a sample in which this RNA was not reverse transcribed to cDNA was used as the negative (“–”) control. To perform gene expression analysis, specific primer sets were designed to produce 100 to 200 bp PCR products (Table S1). Reverse transcription PCR (RT-PCR) was performed with 2×Hieff™ PCR Master Mix (With Dye) (#10102ES08, YEASEN, China), using *ppc* and *secA* as references.

### Real-Time PCR analysis

Real-time PCR analysis was performed as described(Zheng et al. 2020; Hu et al. 2017) with some modifications. Specific primers were designed to produce 100 to 200bp PCR products (Table S1) using *ppc* and *secA* as references. Quantitative realtime PCR was performed using the ChamQ Universal SYBR qPCR Master Mix (# Q711-02, Vazyme, China) by a Bio-Rad CFX96 Thermal Cycler. Differences in expression were calculated by melt-curve analysis using Bio-Rad CFX manager software v3.0 (Bio-Rad). The PCR conditions were as follows, 95°C for 1 min followed by 40 cycles of 95°C for 5 s, 55°C for 30 s. The melting curve was completed by 65–95°C and 5 s increased by 0.5°C.

### Extraction of microcystin

The 7942M strain was cultured at 30°C to the stationary phase. The intracellular and extracellular MCs were extracted separately. The extracellular metabolites were directly enriched using a solid phase extraction (SPE) cartridge (Waters, Sep-Pak Classic C18 Cartridges), washed with 20% (v/v) methanol to remove impurities, and eluted with 100% methanol. The methanol was evaporated to dryness in a rotary evaporator (180 rpm, 38°C, pressure dropped from 300 to 95 mbar) (Innoteg-ScienceOne Vap Basic and IKA VC10), and the sample was resuspended in 1 mL 50% methanol (v/v) in water for LC-MS/MS or 1 mL MilliQ water for enzyme-linked immunosorbent assay (ELISA). To extract intracellular MC-LR, the cultures were treated with 5% aqueous acetic acid (v/v) to destroy the cell walls, and intracellular microcystins were extracted using 80% methanol (v/v). The MCs were enriched using an SPE cartridge, washed with 20% methanol in water to remove impurities, and eluted with pure methanol. The sample was dried in a rotary evaporator and resuspended in 1 mL 50% methanol in water for LC-MS/MS or 1 mL MilliQ water for ELISA. For all quantitative analyses, MC-LR standard (#101043-37-2, Taiwan Algal Science Inc) samples had gone through all procedures to evaluate the lose rate during operations. All methanol used in this extraction process was High Performance Liquid Chromatography (HPLC) grade methanol.

### Qualitative analysis of MC-LR

LC-MS/MS analysis was performed on a Xevo TQ-XS mass spectrometer coupled to an Acquity UPLC I-Class System (Waters). Ten microliters of sample solution were injected into a Waters CSH C18 RP column (100 × 2.1 mm, particle size of 1.7–μm). Multiple reaction monitoring (MRM) mode was used to confirm the peak time. The precursor and fragment ions were m/z 498.61–135.11 and m/z 498.61–861.27. Chromatography was performed at 40°C using 0.1% formic acid in water (A) against methanol (B) at 300 μL min^-1^ using the following gradient: Time 0=2% B, 2=2% B, 10=80% B, 12=80% B, 13=2% B, 15=2% B. The capillary voltage was 3.5 kV, and the desolvation temperature was 350°C.

Mass scan spectra were acquired in positive ion mode over a range of m/z 450–1000. The capillary voltage was 1.5 kV, and the cone voltage was 8 V. The m/z 498.3 molecule was analyzed by MS2 scan: the molecular weight range of the MS2 scan spectra was m/z 100–400, the capillary voltage was 1.5 kV, the cone voltage was 15 V, and the collision energy was 35 eV Chromatography was performed at 40°C using 0.1% formic acid in water (A) against methanol (B) at 300 μL min^-1^ using the isocratic gradient: Time 0=60% B, 4=60% B.

### MC-LR specific enzyme-linked immunosorbent assay (ELISA)

ELISA was performed using a MC-LR ELISA kit based on the use of the monoclonal antibody anti-MC-LR, in which MC-LR or antigen was adsorbed into the reaction wells. ELISA was carried out according to the manufacturer’s instructions.

### Protein phosphatase 2A (PP2A) inhibition assay

MCs were detected using a Microcystins PP2A Kit (#520032, Abraxis) as described by the manufacturer.

### Production and purification of FtsZ

FtsZ was expressed in Rosetta (DE3) cells using pET-21b vector, with C-terminally His6-tagged. Primers used for the tagged proteins were 5’-cgccatatgatgaccgaccctatgcc −3’ and 5’-ccgctcgaggggtcggttttgaattttccg −3’. Overnight culture in fresh LB medium at 37°C with dilution ratio of 1:1000 (culture volume). At OD600 of 0.6, 0.6 mM isopropyl β-D-thiogalactopyranoside (IPTG) was added in the culture, which then transferred to 16°C for overnight culture. The cells were harvested by centrifugation at 9000 g for 10 min at 4°C, and the pellets were wash once with buffer A (20 mM Tris-HCl (pH 7.9), 150 mM NaCl) and stored at –80°C.

The cell pellets were resuspended in 30 mL of buffer A and broken by an Ultra High Pressure Nano Homogenizer (AH-1500 PLUS). The lysates were centrifuged at 16000 g for 1 h at 4°C. The supernatants were purified using an affinity Ni^2+^ column (Qiagen). All proteins were eluted by imidazole of 8 mM, 20 mM, 30 mM, 50 mM, 100 mM, 150 mM, 250 mM, respectively. The FtsZ protein was eluted by100 mM imidazole. Purified FtsZ protein was concentrated with a centrifugal filter (Amicon), measured by Ultra-Micro Nucleic Acid Protein Analyzer (Thermo Fisher Scientific), and checked by SDS-PAGE analysis.

### Immunofluorescence Microscopy

Immunofluorescence microscopy on FtsZ localization was performed as previously described by Miyagishima et al(Miyagishima, Wolk, and Osteryoung 2005).

### Analysis of MicroScale Thermophoresis (MST)

The GTP and MC-LR affinity of the purified FtsZ from *Synechococcus* 7942 were measured using the Monolith NT.115 (Nanotemper Technologies)(Dong et al. 2018). The FtsZ protein with concentration of 4 μM was labelled using fluorescent dye RED-tris-NTA by mixing and incubating for 30 min at room temperature in the dark. For the MST assay, the initial concentration of 1 mM GTP and 1 mM MC-LR were gradient diluted in 16 different serial tubes using PBS Buffer with 10 μL volume. The 10 μL labelled FtsZ was mixed with the 16 tubes described above in different series at room temperature. For competitive binding assay, 4 μM FtsZ protein mixed with 1 mM MC-LR, labelled using above fluorescent dye under the same conditions, and then mixed with the 16 different serial tubes of GTP. The mixed samples were loaded into a high-quality capillaries (NanoTemper Technologies) and measured at room temperature by using 20% LED power and medium MST power. Each assay was repeated 3 times. The data analyses were performed by an analysis software (MO.Affinity Analysis v.2.2.4).

### 90° angle Light scattering assays of FtsZ polymerization

90° angle light scattering for FtsZ polymerization was measured by a F-4700 Fluorescence Spectrophotometer (Hitachi) (Mukherjee and Lutkenhaus 1999; Olson, Wang, and Osteryoung 2010). Both excitation and emission wavelengths were set at 350 nm, with the slit width of 1 nm in the assays. 12.5 μM FtsZ was incubated in PBS buffer, and then placed in the 1 cm path cuvette chamber. Data were collected for 8 min to establish a baseline. 1 mM GTP, 1mM MC-LR and the mix of 1 mM GTP and 1mM MC-LR were mixed respectively with above FtsZ, and the data were collected for 60 min.

### Statistical Analysis

Three biological replicates were performed for each experiment. All data were analyzed using SPSS ver. 19. A *t*-test was applied to evaluate the means and SD of the replicates. The differences between the control and test values were measured by oneway ANOVA, and significant differences were determined as *P* < 0.05 or *P* < 0.01.

## Abbreviations

MCs: microcystins
MC-LR: Microcystin-LR
TAR cloning: transformation-associated recombination
*mcy* cluster: microcystin biosynthetic gene cluster
PKS: polyketide synthase
NRPS: nonribosomal peptide synthetase
7942M: 7942-NSII-mcy cluster
LC-MS: liquid chromatography-mass spectrometry
MS: mass spectrometry
WT: wildtype *Synechococcus* 7942
MST: MicroScale Thermophoresis

## Author Contributions

Q.W. designed the research; Y.Z., C.X., H.C., A.J., L.Z., and J.Z. carried out the experiments; Y.Z., H.C., L.Z. and Q.W. analysed the data; Y.Z., H.C., and Q.W. wrote the manuscript. All authors discussed and commented on the results and the manuscript.

## Data and Materials Availability

The *mcy* gene sequence and NSII sequence information are available on GenBank with accession codes AF183408.1 (https://www.ncbi.nlm.nih.gov/nuccore/AF183408.1) and CP000100.1 (https://www.ncbi.nlm.nih.gov/nuccore/CP000100.1?report=genbank&from=81217&to=81918).

## Supplementary Materials includes

Fig. S1 to S10

Table S1

