## Supplementary materials for "Reconstitution and Expression of *mcy* Gene Cluster in The Model Cyanobacterium *Synechococcus* 7942 Reveals a Roll of MC-LR in Cell Division"

**The file includes:**

Fig. S1 to S10

Table S1

### Figures

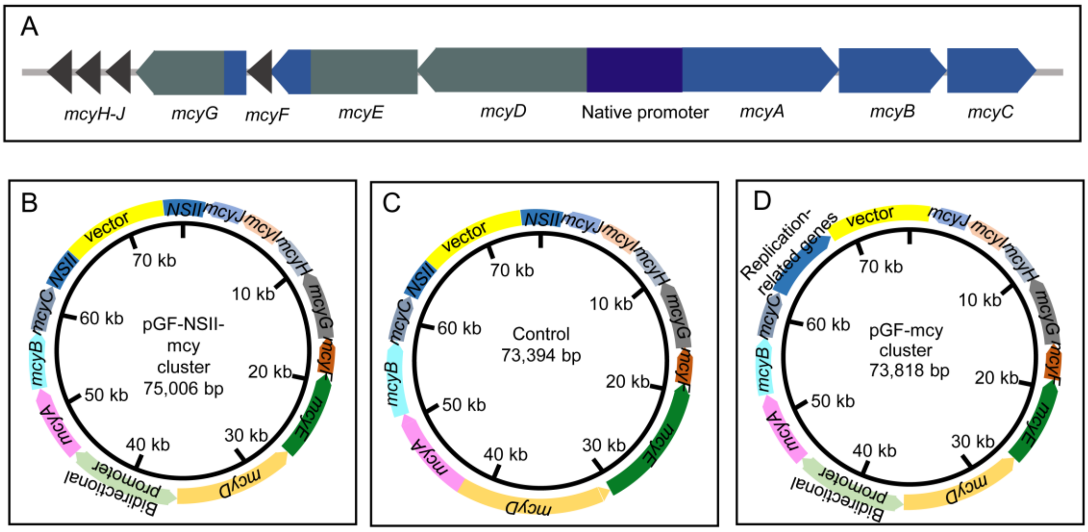


**Figure S1. The mcy gene distributions and the plasmid profile.**

(A) Gene distributions in the MC biosynthesis gene cluster. (B) The pGF-NSII-mcy cluster. (C) No-promoter plasmid control. (D) The pGF-mcy cluster.

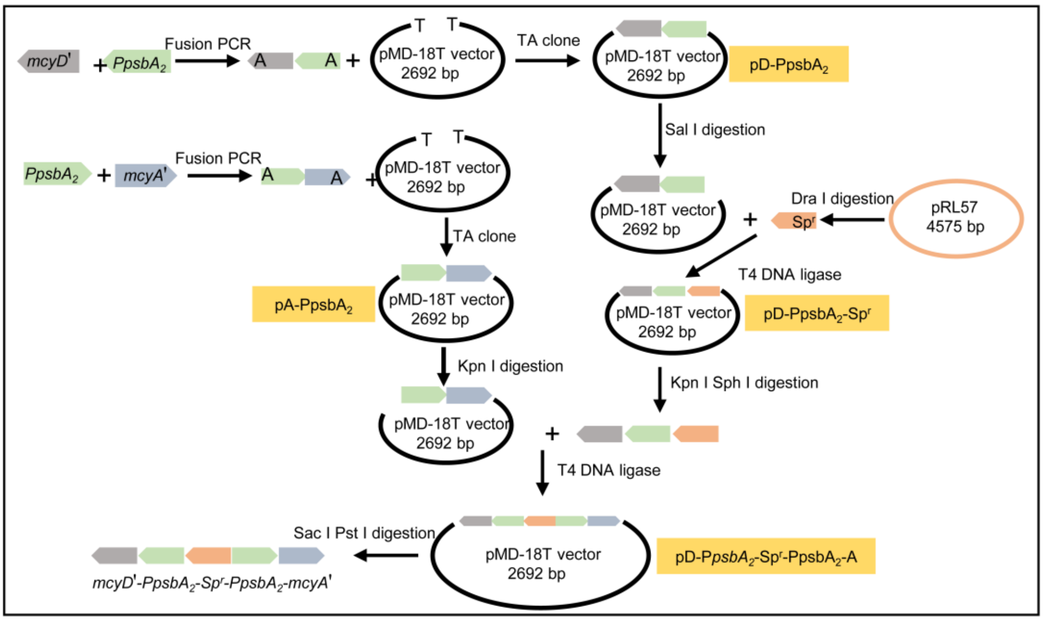


**Figure S2. Construction of the *biPpsbA$_2$* promoter.**

*mcyD'* and *mcyA'* (from *mcyD* and *mcyA*) were recombined with *PpsbA$_2$* by fusion PCR and inserted into the linearized pMD-18T vector by TA cloning to generate D-PpsbA$_2$ (pD-PpsbA$_2$) and A-PpsbA$_2$ (pA-PpsbA$_2$), respectively. pRL57 was digested with Dra I to generate the *Sp$^r$* gene fragment. pD-PpsbA$_2$ was linearized using Sal I, and the *Sp$^r$* gene was inserted into linearized pD-PpsbA$_2$ to generate D-PpsbA$_2$-Sp$^r$ (pD-PpsbA$_2$-Sp$^r$). pD-PpsbA$_2$-Sp$^r$ was digested with Kpn I and Sph I to produce D-PpsbA$_2$-Sp$^r$. pA-PpsbA$_2$ was linearized using Kpn I, and the D- PpsbA$_2$-Sp$^r$ fragment was inserted into linearized pA-PpsbA$_2$ to generate D-PpsbA$_2$-Sp$^r$-PpsbA$_2$-A (pD-PpsbA$_2$-Sp$^r$-PpsbA$_2$-A). Finally, pD-PpsbA$_2$-Sp$^r$-PpsbA$_2$-A was digested with Sac I and Pst I, producing the fragment mcyD'-PpsbA$_2$-Spr-PpsbA$_2$-mcyA', which was named A15. Gray, *mcyD'*. Green, *PpsbA$_2$*. Blue, *mcyA'*. Orange, *Sp$^r$*. Yellow, intermediate plasmids pD-PpsbA$_2$, pA-PpsbA$_2$, pD-PpsbA$_2$-Sp$^r$, and pD-PpsbA$_2$-Sp$^r$-PpsbA$_2$-A.

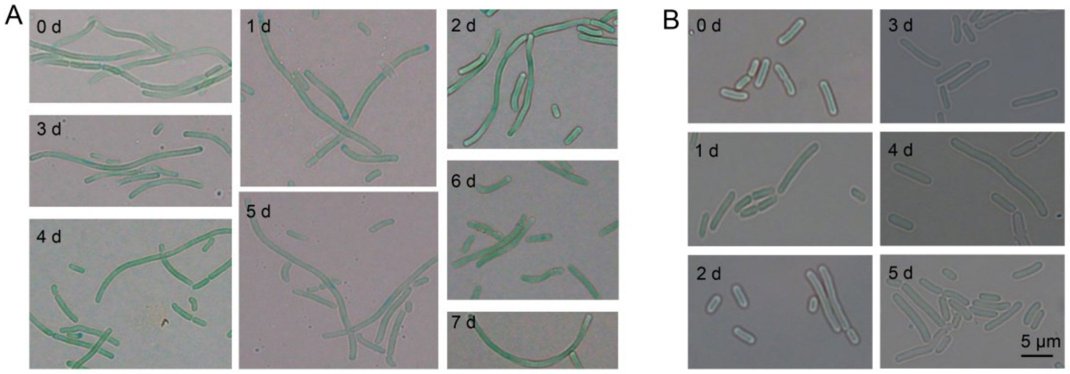


**Figure S3. Microscopic images analysis.**

(A) Daily microscopic images of 7942M cells. (B) WT7942 fed with MC-LR by spreading 20 μg MC-LR on a 70 mm petri dish.

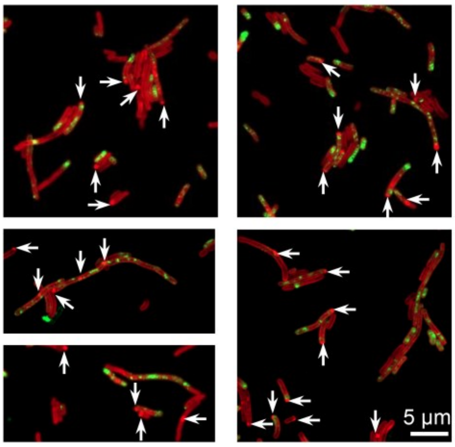


**Figure S4. Localization of FtsZ in 7942M.**

Arrows indicate polarized aggregation of phycobilisomes.

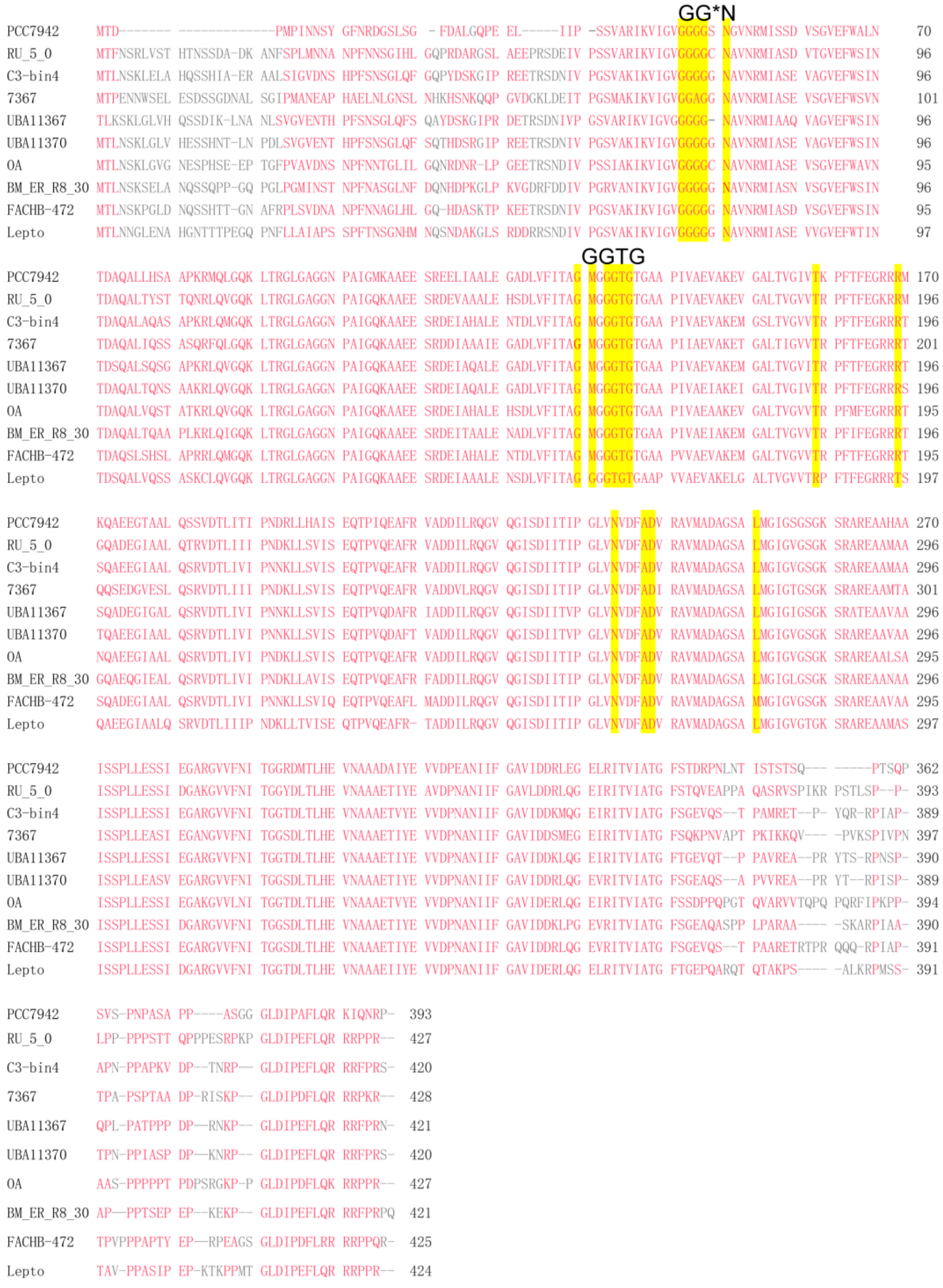


**Figure S5. Sequence alignment of FtsZ from different cyanobacteria species.**

The predicted NBDs are highlighted in yellow, the GG*N (47-51) and GGTP (133-136) domain are marked above; Red letters showing conserved amino acid residues.

Abbreviations: PCC7942, *Synechococcus elongatus* PCC7942; RU_5_0, Cyanobacteria RU_5_0; C3-bin4, *Coleofasciculus* sp.C3-bin4; 7367, *Pseudanabaena* sp. PCC 7367; UBA11367, Cyanobacteria UBA11367; UBA11370, Cyanobacteria

UBA11370; OA, *Oscillatoria acuminate*; BM_ER_R8_30, *Chroococcidiopsidaceae* BM_ER_R8_30; FACHB-472, Cyanobacteria FACHB-472; *Lepto, Leptolyngbya sp*.

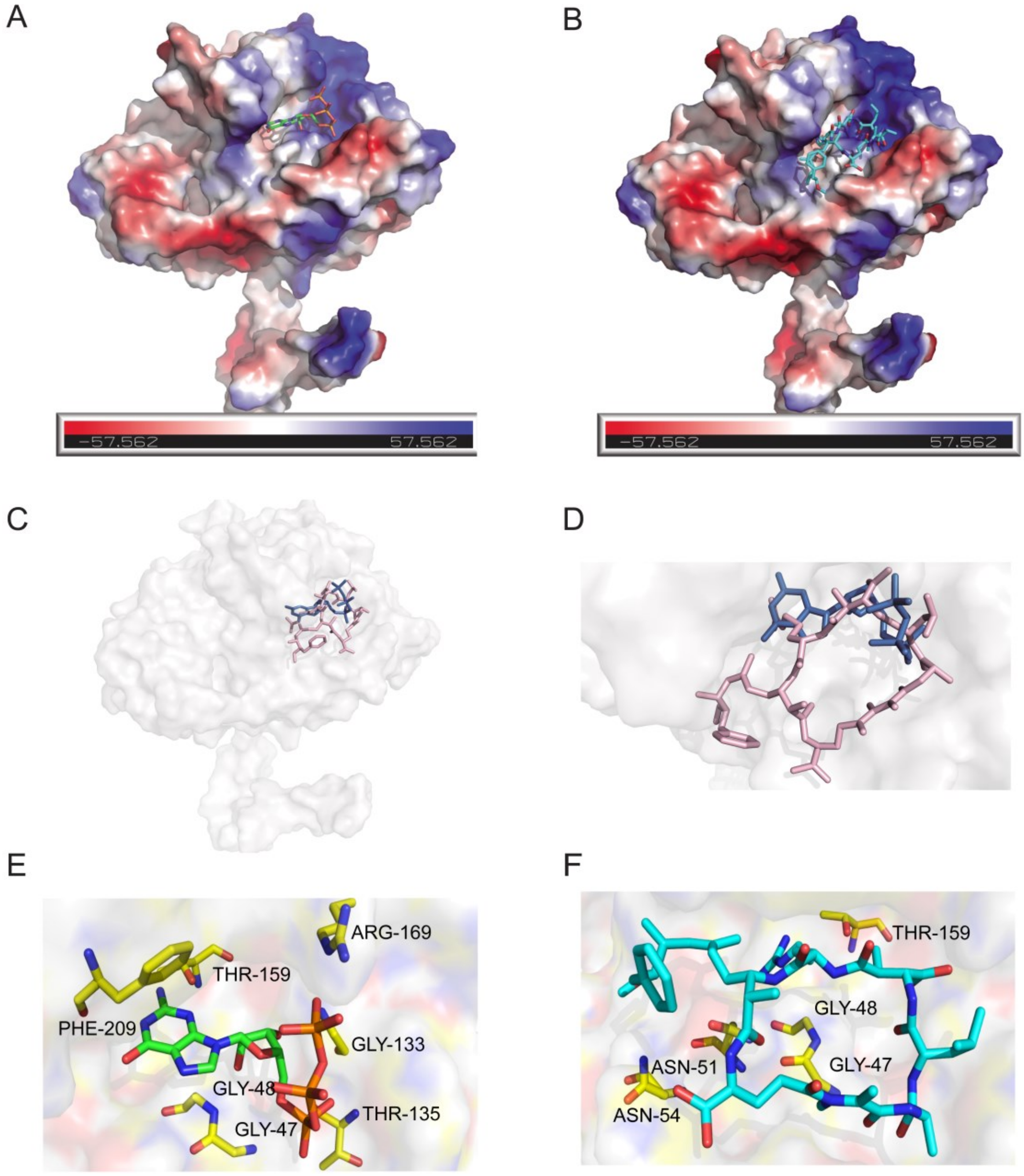


**Figure S6. Molecular docking analysis for FtsZ monomer with GTP and MC-LR.** (A) and (B) Electrostatic potential map of FtsZ monomer docking with GTP (docking score –7.1) and MC-LR (docking score –8.3), respectively. (C) and (D) General and close view of both GTP (blue) and MC-LR (pink) compete for the same binding pocket of FtsZ monomer. (E) and (F) Interactive amino acid residues of FtsZ monomer to GTP (GLY47, GLY48, GLY133, THR135, THR159, ARG169 and PHE209) and MC-LR (GLY47, GLY48, ASN51, ASN54 and THR159).

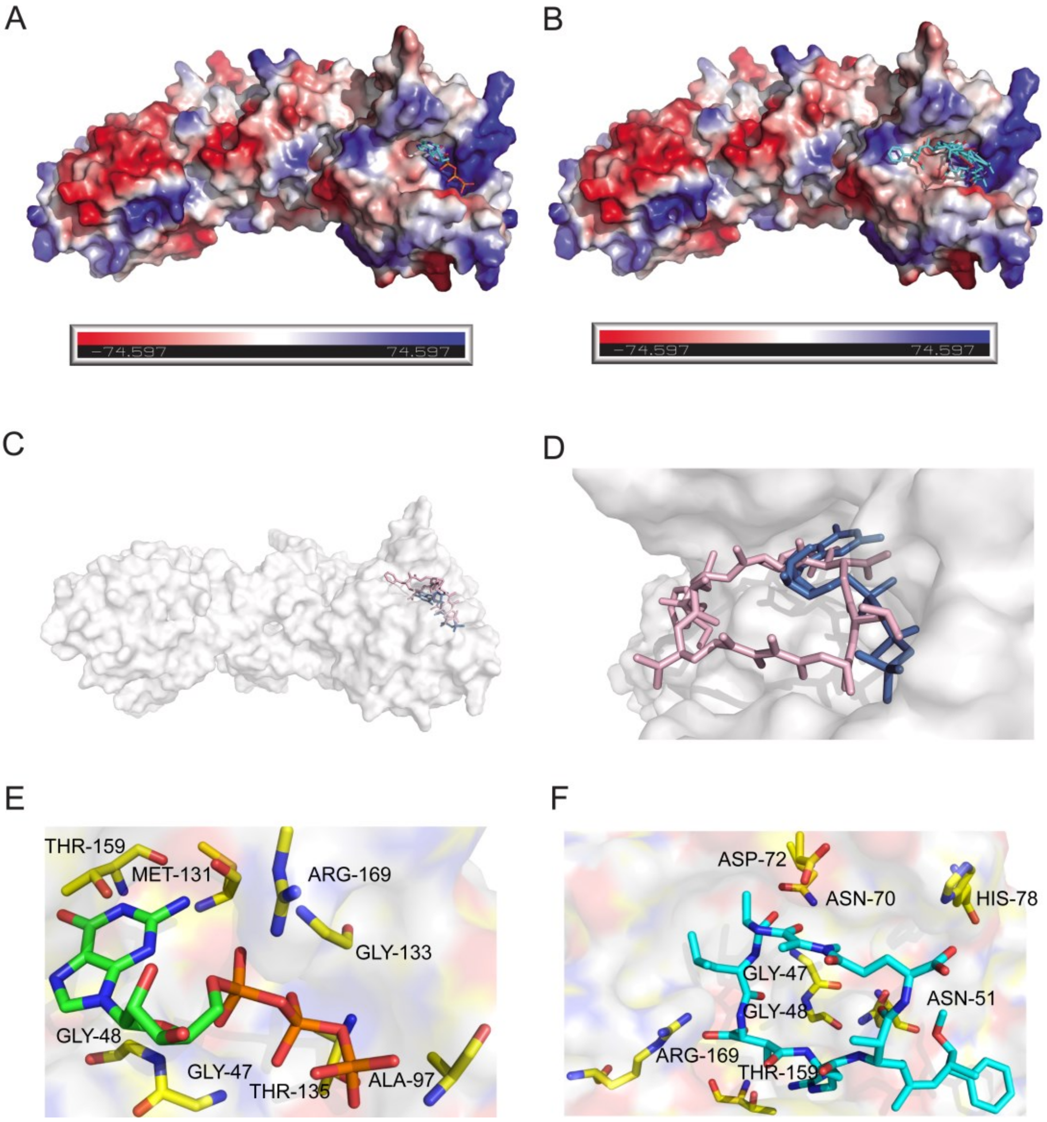


**Figure S7. Molecular docking analysis for FtsZ dimer with GTP and MC-LR.**

(A) and (B) Electrostatic potential map of FtsZ dimer docking with GTP (docking score –9.1) and MC-LR (docking score –9.3), respectively. (C) and (D) General and close view of both GTP (blue) and MC-LR (pink) compete for the same binding pocket of FtsZ dimer. (E) and (F) Interactive amino acid residues of FtsZ dimer to GTP (GLY47, GLY48, ALA97, MET131, GLY133, THR135, THR159 and ARG169) and MC-LR (GLY47, GLY48, ASN51, ASN70, ASP72, HIS78, THR159 and ARG169).

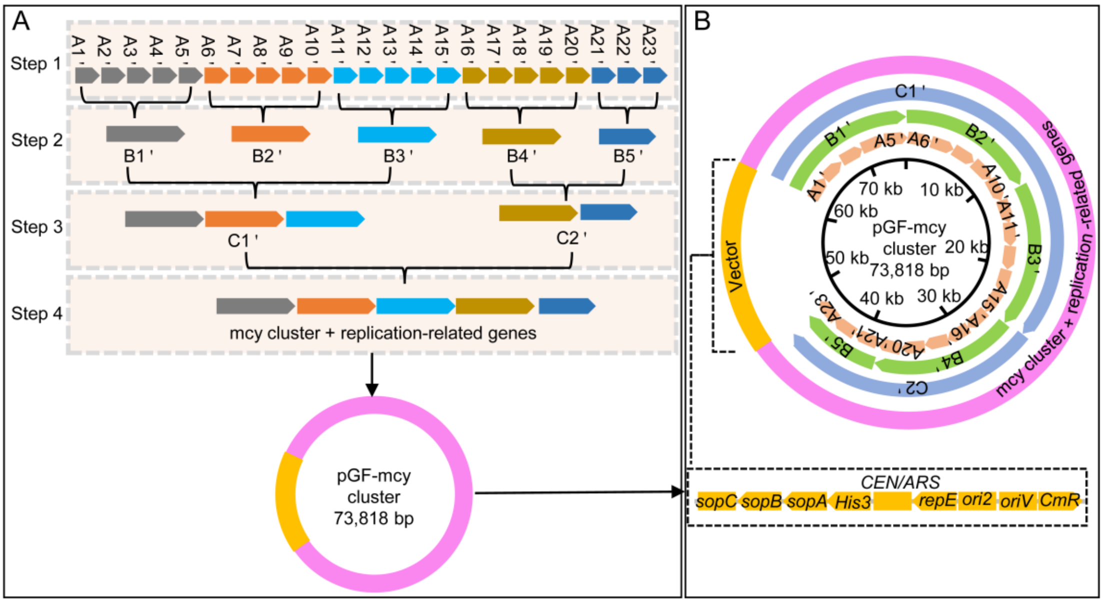


**Figure S8. The process used to assemble the autonomously replicating plasmid pGF-mcy cluster in vitro (similar to the assembly of the pGF-NSII-mcy cluster).** (A) The mcy cluster and replication-related genes were divided into 23 A' fragments, A13' is *biPpsbA*$_2$; intermediate plasmids B' and C' were assembled using 3–5 A' fragments and 2–3 B' fragments respectively; the pGF-mcy cluster was synthesized using two C' fragments. Gray arrows, A1'–A5' and its recombination sequence. Orange arrows, A6'–A10' and its recombination sequence. Blue arrows, A11'–A15' and its recombination sequence. Brown arrows, A16'–A20' and its recombination sequence. Dark blue arrows, A21'–A23' and its recombination sequence. (B) A circular map of the mcy cluster and the corresponding position of the mcy cluster and TAR vector region. The light orange arrows represent 23 A' fragments in the inner circle; the green arrows represent five B' fragments in the second layer from the inside; the blue arrows represent two C' fragments in the third layer from the inside; purple represents the mcy cluster, *biPpsbA*$_2$ and replication-related genes; yellow represents the vector region.

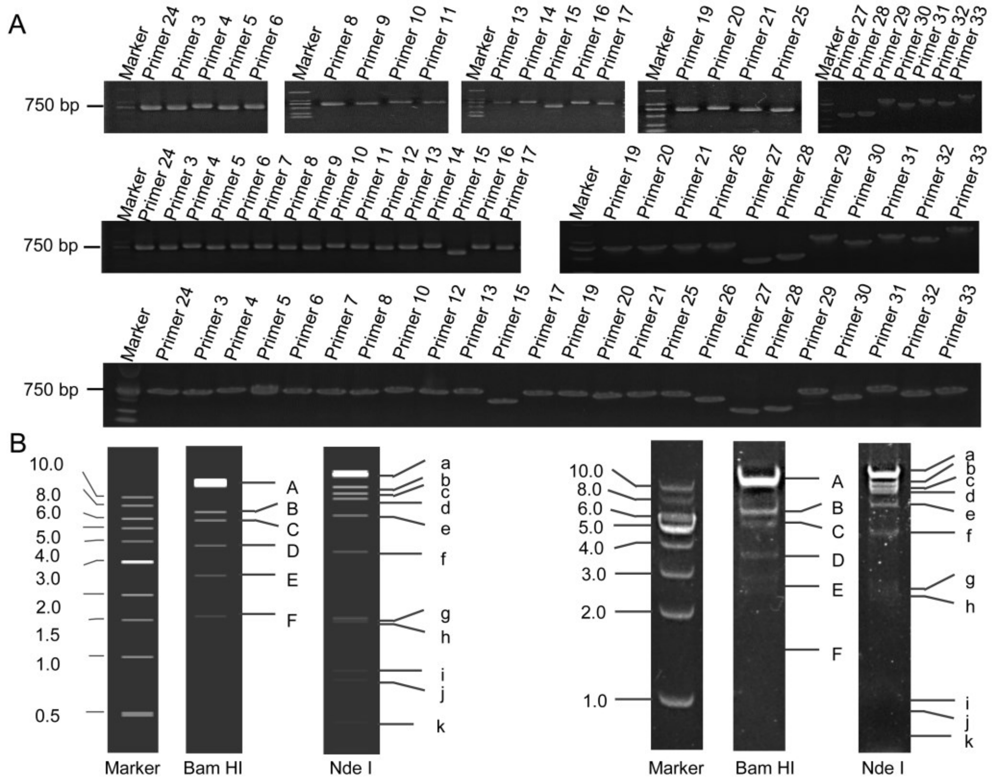


**Figure S9. The verification used to assemble the autonomously replicating plasmid pGF-mcy cluster and the intermediate plasmids in vitro.**

(A) PCR analysis of the intermediate plasmids B' (upper) and C' (middle) and pGF-mcy cluster (lower). (B) The restriction enzyme profiles of pGF-mcy cluster predicted by SnapGene 4.1.9 software (left) and the actual restriction enzyme digestion profiles of pGF-mcy cluster following 1% agarose gel electrophoresis (right).

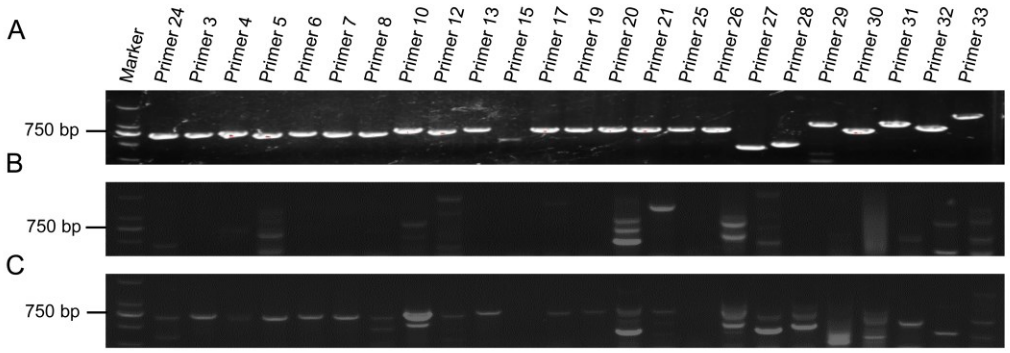


**Figure S10. PCR analysis of the 7942-mcy cluster transformant.**

(A) The genome of 7942-mcy cluster from a first generation transformant was used as a template. (B) The genome of 7942-mcy cluster from a fifth generation transformant was used as a template. (C) The genome of 7942-mcy cluster from a reactivated first-generation transformant from an ultra-low temperature freezer was used as a template.

### Tables

**Table S1.** Primers used in this study.

| Name | Sequence (5'⟶3') |
|---|---|
| McyD'-F | TGAGGTAAGAATACGGGCATAGT |
| McyD'-R | AATTATAACCATGGACTTTCAAGATAAAAAGAACTTAT |
| McyD'-PsbA$_2$-F | GAAAGTCCATGGTTATAATTCCTTATGTATTTGTCG |
| McyD'-PsbA$_2$-R | TGCGGCTTTAGCGTTCCAGTGGAT |
| McyA'-F | TGCGGCTTTAGCGTTCCAGTGGAT |
| McyA'-R | CAATGCTTTTGGTTATAATTCCTTATGTATTTGTCG |
| McyA'-PsbA$_2$-F | AATTATAACCAAAAGCATTGTACCCCATGAC |
| McyA'-PsbA$_2$-R | CCCATTTTCTGAAGTCGTCAAG |
| Primer1-F | GTTCCGCCCGACTGAGAT |
| Primer1-R | CGCAGATAAGGCCAAAGG |
| Primer2-F | TGCCCGATGGCTATGACA |
| Primer2-R | TGAATAATCTGCAAATTGCACTC |
| Primer3-F | GTTCTAACACCCATTGATGATACTT |
| Primer3-R | AATCCTTGAATTTGGTTCTTTG |
| Primer4-F | ACAGGAAGCGGACTGGTAATG |
| Primer4-R | CTGGCTAATTTTTACTGATGTTGAAC |
| Primer5-F | AGGGATTTACCCGTTAGTTTTTG |
| Primer5-R | CATATGCAGATTGCTTCCGG |
| Primer6-F | TGAATGAGTTCTAACCATTGCAG |
| Primer6-R | AACTGCCACTAAACATAATAATCAT |
| Primer7-F | ATTATGGCTGATTGATTCTTCG |
| Primer7-R | CCAGTTTATATGATTCCGAGTTAC |
| Primer8-F | AACTGCATCGGTGGTGTGAG |
| Primer8-R | TGTTTTCTCTCCACCTGCCAT |
| Primer9-F | TCTGCTAAATTTTCGCCGC |
| Primer9-R | ACAAACCTCTAACCTTACCGATACC |
| Primer10-F | CCCCTGATGGACGGCTAC |
| Primer10-R | GGATAGGTAGATGAACATGAGTAAGC |
| Primer11-F | ATATAACTCTTGAATCGCTTGGAAC |
| Primer11-R | CTTATTTCCGCTAACCAAGATCTAT |
| Primer12-F | CTAGGAATAATGCCAAAAAATTCTC |
| Primer12-R | CTGGTGTAATTAAGATTGACTGGC |
| Primer13-F | ATTAAGCGCTCATATTGCGC |
| Primer13-R | GAAGACACATTACAGTTAATCATTG |
| Primer14-F | CCAAACCCATCGGCACTT |

| | |
|---|---|
| Primer14-R | GAAAACGACCAATCAATCCAA |
| Primer15-F | TGCGGCTTTAGCGTTCCAGTGGAT |
| Primer15-R | GGTTATAATTCCTTATGTATTTGTCG |
| Primer16-F | GCGGCGATTGATGTGACT |
| Primer16-R | CCACAACCTATTTCCAGAACTTT |
| Primer17-F | TCAAGATTGGGATTGGTTAGTTC |
| Primer17-R | GGATAGTGAACCTCAGTATCATT |
| Primer18-F | GAGAATGTAGAACTTCCACCTAAAAC |
| Primer18-R | TCTGGCCGTAATCTCTAATAAATG |
| Primer19-F | AACAGCAGGTGCGAGTCTTG |
| Primer19-R | TGCTGTAAGAAACGGCGC |
| Primer20-F | CAATTAAGCAAACTGCAACCC |
| Primer20-R | CATTCAAAGGATTAGGCACAAA |
| Primer21-F | TGGGGATTTTTATTGAGCGT |
| Primer21-R | GGGAATTGGGCTGACAGCAC |
| Primer22-F | CTGAATAATCTGCAAATTGCACT |
| Primer22-R | CCCGTGACGGGCTACACC |
| Primer23-F | AGCACCGACGCAAGGTCA |
| Primer23-R | GGCAACCGTCTATCCCACC |
| Primer24-F | TGTCGGGGCTGGCTTAAC |
| Primer24-R | AGATTCGGAGAATGAATTAGCC |
| Primer25-F | CAAGGGACTATTCATTTGATTCG |
| Primer25-R | GGGTTTGCAGGTTTGGGG |
| Primer26-F | CCCCCACCAGCACCTTG |
| Primer26-R | AACGCCTGGTTGCTACGC |
| Primer27-F | CCGATCCTCCGGCCACTC |
| Primer27-R | TGCTTTTGCTTTTCGGCTCC |
| Primer28-F | CCGATGAAGTGGTGGAGCA |
| Primer28-R | GGTCAGTCCTCCATAAACATTG |
| Primer29-F | ATGGCGATTTATCACCTTACGG |
| Primer29-R | GCCATCTCGCTGCGGTACT |
| Primer30-F | CAGGGCAATGACGTGTATATCA |
| Primer30-R | AAATGTCTTGTTTACAATAGAGTGGG |
| Primer31-F | ATGAAGAACGACAGGACTTTGC |
| Primer31-R | CTACATGCTGAAATCTGGCCC |
| Primer32-F | ATGGCTACCCATAAGCCTATCA |
| Primer32-R | TTAGGCTTCACCACGGGG |
| Primer33-F | CGCTCAAGCACATACAGGACT |
| Primer33-R | CAGGAAACAGCTATGAC |
| mcyA-F | GAACCAGTTTCGCACTCTTTG |
| mcyA-R | TCAATAGCCGTGTCTTGTTCAG |

| | |
|---|---|
| mcyB-F | ATTGAACGGATGGCAGGAC |
| mcyB-R | GCCTCCCTATTGTTCCACTCT |
| mcyC-F | TATGATCCCTTCTGCCTTTGT |
| mcyC-R | CGGGGTGCTAAATAGGGTG |
| mcyD-F | CCGGAACCTTAGACGATACAA |
| mcyD-R | TCTAATGGGTCATTTAGGGTGA |
| mcyE-F | ACCTACACTGGCGGACAAGA |
| mcyE-R | GCGGCGAACATCGTCATA |
| mcyF-F | TTGCATAGTCATTGGCTGTTG |
| mcyF-R | GGAGTTCTTGGTCCGCTATTT |
| mcyG-F | ATGCCCTTGGATCATGTAGG |
| mcyG-R | GCAGAGGTTGGCGTAGGAC |
| mcyH-F | TTTAGTCGAGGTTATCGCTCAG |
| mcyH-R | CTGCAAGCAATAATAGCCTGAT |
| mcyI-F | ATTGGTACTCTGGTGGCTCAC |
| mcyI-R | GCTCCTACTGTCTCCGCTTT |
| mcyJ-F | GCCCAGATTAGCCAAGGAA |
| mcyJ-R | AAACGCCATAAACCAACCC |
| 0080-F | GCGGTCTACTTGCCCTTGA |
| 0080-R | TTGGTGATTTGAAACTGTGCC |
| 0081-F | CAGTTTCTGAAATGGATGGGAG |
| 0081-R | GGCCTCGGCGAGTTTCTAC |
| 0082-F | CGGAACCCTACGAAATCGA |
| 0082-R | CCCAGCACGACAATGAAGA |
| 0083-F | GACGCCCCAGACTGAATTG |
| 0083-R | GCAGCAAGCGTTTTCCATAC |
| 0085-F | CATCTACAAGCCGTTCCTGG |
| 0085-R | CGCTGAAGATTCGTCCCTC |
| 0086-F | TTGATGAGTTCCGCGACCT |
| 0086-R | TTCGCTAGACCTTTGAAAGACAG |
| ftsZ-F | GGGCTGACCTCGTCTTTATC |
| ftsZ-R | GCAGTGCGGCTGTTCCT |
| minC-F | CCTGGGTGATTGGCTGTTC |
| minC-R | GGGCGTCAGTTGGAATGG |
| minD-F | GCAACCTGAACCGCCACT |
| minD-R | GCATCAACGAGGAAACTCTGC |
| minE-F | GAGCTGGACTCTGAGGGAATG |
| minE-R | CAGCAGTTGCGGGTTTGAC |
| zipN-F | GGGCTGTCCGAGCAATAGG |
| zipN-R | GCCTGCTACACCCTGATTGC |
| sulA-F | GAAGAACGGGCGCTAACTG |

| | |
|---|---|
| sulA-R | CCGGCAACACGGTCTAATAC |
| ppc-F | CGGGACTGAGCTGTTACGA |
| ppc-R | TGGAGGTATTCACAGATTTCGT |
| secA-F | CGTCACCGTCAACGACTACTT |
| secA-R | CGACATTCCCTGCTGGATTAG |
| A1-F | TCGATCATGAACTGATTGAGGAT |
| A1-R | CAGCACGACAATGAAGAGAATG |
| A2-F | ATCCTGCAGCTGATCGTCG |
| A2-R | CTGAATAATCTGCAAATTGCACT |
| A3-F | TCTTGCCTAATGCTTTATCCG |
| A3-R | GAACAATTTCTCTTTTTACCTCAAC |
| A4-F | AGTTCTCCATCGGTCATTTCAC |
| A4-R | CCTAATCCCCCTCAATTATTCC |
| A5-F | ATTTTCTAGGGCAATAACACGC |
| A5-R | CATTTGCTCACCCTTTCCG |
| A6-F | CAGATCCTTTTCAGGTTGACTTTC |
| A6-R | TATGATCTGGAAAATTCTGCCG |
| A7-F | TTTCAGTAATTAGCTCACTGTCGG |
| A7-R | TAAATGTATCCGTAAAATTGGCTG |
| A8-F | TTGATAATTGTATTGAGTCTGGGAG |
| A8-R | AGCATCCTATAACGTCACAACATC |
| A9-F | GCTTCATGGCGATTAATAACTTG |
| A9-R | TGTCAAAGAAAAACCCTTAGAAAATAC |
| A10-F | CCTTGATGATATAGGCTGTAGCG |
| A10-R | GGGATGGCTTGTCGTTTTC |
| A11-F | TGGTTTTTGAGAATACAGTTTTGAG |
| A11-R | ACATGATCTTTGAAGGTGAGGTG |
| A12-F | GACTTGCGGCTAAAATGGC |
| A12-R | TTTGGGGATGGACTCTCTCAC |
| A13-F | GCTTCAACATTCGGAAAACG |
| A13-R | AAATCAGGGATAAATTACGGGAG |
| A14-F | TTTAAATAAAATGGCAGCAATCC |
| A14-R | TTAGCTATCGTGGGTTTAGGTTG |
| A15-F | TGAGGTAAGAATACGGGCATAGT |
| A15-R | CCCATTTTCTGAAGTCGTCAAG |

| | |
|---|---|
| A16-F | AAGTTAAGATTCGAGGTTTCCG |
| A16-R | CTCCATCTGCTTTTCCCCTAG |
| A17-F | ATCATCGGGGTGAAAGATTTAG |
| A17-R | AGCATCATCACTCAAATATCGC |
| A18-F | AGGACGGTGGGTTGGTTTACA |
| A18-R | AAGATAGCGAACTAAATCCCCAG |
| A19-F | TTTTAGATCAGAATTTACAACCGC |
| A19-R | CCATTCAGCCAGTTCTTTCAC |
| A20-F | CACACTGTCCCCTCGGTAATG |
| A20-R | GAATATAAACTTGGACATTAGCAATGG |
| A21-F | AGATCGCATTCTCCAAATTACTC |
| A21-R | TCTTGCCTAATGCTTTATCCG |
| A22-F | CTGAATAATCTGCAAATTGCACT |
| A22-R | GAATCTTGAAGCTTGTCATCTGC |
| A23-F | CCCGTGACGGGCTACACC |
| A23-R | GAATCTTGAAGCTTGTCATCTGC |
| pGF-F | AGTCACTGCCAGGTATCGTTTG |
| pGF-R | ACGTTAATCACTTGCGATTGTGT |
| pGF-A1-R | CGCACCCAAGAAATTGGCCTGCGCAAAGCGATCGGGGCGACC<br>CAAAAAGACATCCTCAATCAGTTCATGATCGAACGCGTCGGGT<br>ACCGAGCTCGAATTC |
| pGF-A7-F | AGTTATCAAAAATACCGATAGGATGTTTAGAGAGAATTTTTTCC<br>CAGATACTAACGAGATTGGATTCTAAATAATTCACGCGTCTAGA<br>GTCGACCTGCAG |
| pGF-A8-R | ATTGCTGGATTGGCTACTTTAATCTCCCAGACTCAATACAATTA<br>TCAAACGCGTCGGGTACCGAGCTCGAATTC |
| pGF-A12-F | TAACAGATTTCGTAACTCTGTAGAAGTGAGAGAGTCCATCCCC<br>AAAACGCGTCTAGAGTCGACCTGCAG |
| pGF-A13-R | TTGTTCTCTCCCTTCGACAATTACTTTTCGTTTTCCGAATGTTG<br>AAGCACGCGTCGGGTACCGAGCTCGAATTC |
| pGF-A17-F | CAGGGGAATTGGCTATGGAATTTCGCGATATTTGAGTGATGATG<br>CTACGCGTCTAGAGTCGACCTGCAG |
| pGF-A18-R | GAGTCGCAAAATAACCGGAAAAAGACTGGTAAACCAACCCAC<br>CGTCCTACGCGTCGGGTACCGAGCTCGAATTC |
| pGF-A23-F | CGCCGTTAGCCACTGCTGCAAGACTGGTGGCAAGGGCAAAAC<br>AATCAAAATGAGGATAAAGAGGGCCAAGAAGCCAAACGCGTC<br>TAGAGTCGACCTGCAG |

| | |
|---|---|
| pGF-A1'-R | AGAGAATTAATCCATCTTCGATAGAGGAATTATGGGGGAAGAA<br>CCTGTGCCGGCGGATAAAGCATTAGGCAAGAACGCGTCGGGT<br>ACCGAGCTCGAATTC |
| pGF-A20'-F | GGTGCGCTCTCCGAGGGCCATTGCATGGAGCCGAAAAGCAAA<br>AGCAACGCGTCTAGAGTCGACCTGCAG |
| pGF-A21'-R | AACAAGGCGCAAGGTGCTGGTGGGGGCCATGATTTTGGCCAA<br>GGTGAACAGCAGCGAGTGGCCGGAGGATCGGACGCGTCGGG<br>TACCGAGCTCGAATTC |
| pGF-A23'-F | GCGGCGGGCAAGTACGACATCACCCGGCCCAAGGCGGCAGG<br>CTGACCCCCCCCACTCTATTGTAAACAAGACATTTTACGCGTC<br>TAGAGTCGACCTGCAG |
| A1'-F | TCTTGCCTAATGCTTTATCCG |
| A1'-R | GAACAATTTCTCTTTTTACCTCAAC |
| A2'-F | AGTTCTCCATCGGTCATTTCAC |
| A2'-R | CCTAATCCCCCTCAATTATTCC |
| A3'-F | ATTTTCTAGGGCAATAACACGC |
| A3'-R | CATTTGCTCACCCTTTCCG |
| A4'-F | CAGATCCTTTTCAGGTTGACTTTC |
| A4'-R | TATGATCTGGAAAATTCTGCCG |
| A5'-F | TTTCAGTAATTAGCTCACTGTCGG |
| A5'-R | TAAATGTATCCGTAAAATTGGCTG |
| A6'-F | TTGATAATTGTATTGAGTCTGGGAG |
| A6'-R | AGCATCCTATAACGTCACAACATC |
| A7'-F | GCTTCATGGCGATTAATAACTTG |
| A7'-R | TGTCAAAGAAAAACCCTTAGAAAATAC |
| A8'-F | CCTTGATGATATAGGCTGTAGCG |
| A8'-R | GGGATGGCTTGTCGTTTTC |
| A9'-F | TGGTTTTTGAGAATACAGTTTTGAG |
| A9'-R | ACATGATCTTTGAAGGTGAGGTG |
| A10'-F | GACTTGCGGCTAAAATGGC |
| A10'-R | TTTGGGGATGGACTCTCTCAC |
| A11'-F | GCTTCAACATTCGGAAAACG |
| A11'-R | AAATCAGGGATAAATTACGGGAG |
| A12'-F | TTTAAATAAAATGGCAGCAATCC |
| A12'-R | TTAGCTATCGTGGGTTTAGGTTG |
| A13'-F | TGAGGTAAGAATACGGGCATAGT |
| A13'-R | CCCATTTTCTGAAGTCGTCAAG |

| | |
|---|---|
| A14'-F | AAGTTAAGATTCGAGGTTTCCG |
| A14'-R | CTCCATCTGCTTTTCCCCTAG |
| A15'-F | ATCATCGGGGTGAAAGATTTAG |
| A15'-R | AGCATCATCACTCAAATATCGC |
| A16'-F | AGGACGGTGGGTTGGTTTACA |
| A16'-R | AAGATAGCGAACTAAATCCCCAG |
| A17'-F | TTTTAGATCAGAATTTACAACCGC |
| A17'-R | CCATTCAGCCAGTTCTTTCAC |
| A18'-F | CACACTGTCCCCTCGGTAATG |
| A18'-R | GAATATAAACTTGGACATTAGCAATGG |
| A19'-F | AGATCGCATTCTCCAAATTACTC |
| A19'-R | TCTTGCCTAATGCTTTATCCG |
| A20'-F | TTCCGCGTCCTTGCAATAC |
| A20'-R | TGCTTTTGCTTTTCGGCTCC |
| A21'-F | CCGATCCTCCGGCCACTC |
| A21'-R | GGTCAGTCCTCCATAAACATTG |
| A22'-F | ATGGCGATTTATCACCTTACGG |
| A22'-R | GCCATCTCGCTGCGGTACT |
| A23'-F | CAGGGCAATGACGTGTATATCA |
| A23'-R | AAATGTCTTGTTTACAATAGAGTGGG |
